# Motivational incentives improve working memory performance by enhancing memory-based representations rather than altering response strategies

**DOI:** 10.64898/2026.10.02.755939

**Authors:** Vivek Nair, Owen Agnel, Eda Mızrak, Sanjay G. Manohar

## Abstract

Motivational incentives are known to improve working memory (WM) when items differ in value, but whether they enhance performance when all items are equally valued, and if so, by what mechanism, remains unclear. Improvements could reflect enhanced memory-based representations or speed-accuracy trade-offs, whereby incentives promote slower, more cautious responding without improving memory itself. Here, we tested these alternatives across two experiments using an auditory WM task in which incentives varied across trials, but all items on a trial were equally incentivised. In Experiment 1,we allowed participants to resubmit their responses if desired. Reward-related improvements in WM performance were selectively observed when memory load was high, and only at serial positions associated with greater interference. These improvements were also dissociated from systematic response slowing and resubmission behaviour. This indicated that the benefits of incentives were unlikely to be attributable to response caution. Critically, in Experiment 2, motivational benefits persisted under strict response deadlines, where speed cannot be traded for accuracy. Computational modelling showed that reward selectively improved the precision of memory-based responses without reducing guessing. Together, these findings indicate that motivational incentives improve WM by enhancing the quality of memory-based representations, rather than by strategic slowing or altering response strategies.

## Introduction

Working memory (WM) enables the short-term maintenance and manipulation of information necessary for goal-directed behaviour, but its capacity is fundamentally limited in the number of items that can be held and the precision of recall^1–6^. Despite these constraints, there are ways to improve the quality of internal representations, such as by allocating attention or effortful control^7,8^. Motivational incentives have been shown to influence performance across multiple domains^9–11^, indicating that they can also affect WM representation quality^12–14^. Within WM, such effects are well established when individual memory items differ in value. Incentives improve recall by promoting selective prioritisation during encoding^15–17^: more valuable items are stored with greater precision than less valuable ones^18–22^, incurring a cost to non-prioritised items. This is consistent with incentives redistributing limited memory resources rather than expanding overall storage capacity^2,3,5,23–25^. However, in many real-world situations, all information is equally relevant making prioritisation impossible. Whether motivational incentives can improve WM under such equal-value conditions, and if so, how, remains unresolved.

When all items in WM are equally valued, any motivational improvement must arise from a global enhancement of WM processing rather than selective resource redistribution. Resource-rational accounts propose that WM resource allocation reflects a cost-benefit optimisation process^26,27^: when reward is high, the benefits of improved representational precision outweigh the costs of increased cognitive effort, such that incentives may enhance the quality of memory-based representations even when prioritisation is not possible^7,14,28,29^. However, improved behavioural performance does not by itself prove enhanced memory representation. Reward-related improvements could instead reflect downstream changes during response formation, such as slower or more cautious responding, or post-response refinement, without changing the quality of memory representation. Distinguishing these alternatives is essential for understanding the mechanism by which motivation influences equal-value WM.

A number of equal-value WM studies have shown genuine reward-related improvements in the quality of memory-based representations^30–37^. However, this has been challenged by studies that either do not find the effect^38^ or attribute it to other causes^39^. Van den Berg et al.^38^ used delayed-estimation tasks where participants reproduced the orientation of probed items, manipulating monetary reward both between and within subjects across three experiments. They found no evidence that reward improves WM performance or encoding precision under equal-value conditions, concluding that equal-value visual WM encoding quality may be insensitive to monetary incentives. Conversely, Grogan et al.^39^ reported motivational improvements under equal-value reward conditions, but attributed these to changes to response behaviour rather than enhanced memory-based representation quality. Using continuous-report visual WM tasks involving reproducing the orientation of probed items, they systematically manipulated reward timing and maintenance intervals, finding no evidence that incentives enhanced encoding or maintenance. Instead, because reward was associated with longer response times, they interpreted their findings as a speed-accuracy trade-off (SAT)^40–42^, whereby incentives promote more cautious retrieval-related decision processes^43,44^ without enhancing memory-based representations. Together, these findings leave unresolved whether motivational improvements in equal-value WM can reflect genuinely enhanced memory-based representations or instead arise from changes in response behaviour during retrieval.

These competing accounts make three distinct and testable predictions. First, if incentives enhance the quality of memory-based representation guiding responses, benefits should depend on genuine memory demand (i.e. WM load) and be greater when multiple items must be remembered compared with when memory requirements are minimal, where all items already have very high quality^2–5^. Second, motivational benefits should be most strongly expressed when memory items are subject to the greatest interference, i.e. in situations where boosting the representation strength would confer the greatest advantage^4,45,46^. Third, benefits should persist under strict response time constraints, because higher-quality representations should support more accurate responding regardless of available decision time. They may also be coupled to metacognitive strategies, strengthening the relation between errors and their correction^47,48^. However, if motivational improvements reflect slower, more cautious responding, they should be less tied to WM load and retrieval difficulty (i.e. level of interference), and attenuated when response times are experimentally restricted.

Here, we tested these alternatives using an auditory WM task where all items were equally incentivised. Although auditory WM relies on modality-specific representational systems, it shares core capacity and control constraints with visual WM, providing a useful test of whether motivational modulation of equal-value WM generalises across sensory domains^49–51^. Reward was cued before stimulus presentation, to maximise participants’ ability to adjust resource allocation to the incentive.

In Experiment 1, we manipulated reward and memory load to determine whether motivational benefits depend on WM demand, examining serial-position effects to test whether improvements were greatest where retrieval competition was highest. To test whether motivational effects could be explained by response behaviour changes, participants could resubmit their responses as many times as they wished before final submission, allowing us to examine two response-behaviour markers: response slowing, reflecting response caution, and resubmission rate, reflecting corrective refinement of responses after their initial submission. Trial-by-trial confidence ratings were collected as a metacognitive index of response accuracy: if confidence tracks the quality of memory-based representations, it should predict recall error independently of response time and resubmission. In Experiment 2, we imposed strict response deadlines to directly test whether motivational benefits require extended decision time. Finally, we applied mixture modelling across both experiments to test whether improvements reflected greater precision of memory-based responses or changes in guessing^5,25^.

## Results

Two experiments tested whether motivational improvements in equal-value WM were better explained by enhanced memory-based representations or by changes in response behaviour. In Experiment 1 (**Fig.1a**), participants remembered either one tone (low load) or five tones (high load), maintained the sequence across a 1.5-second retention interval, and reproduced the duration of one of the tones, probed by serial position, by holding down a key. The duration of the keypress constituted the response. Recall error, the primary outcome measure, was computed as the absolute residual from a participant-level regression correcting for central tendency bias (see Methods). The reward level (20p vs 1p) was cued before each trial, and was obtained if the error was below a threshold (see Methods). Participants could resubmit responses as many times as they wished before final submission, and provided confidence ratings after each response. We tested whether reward-related benefits depended on memory load and serial position, and whether they could be explained by slower responses or corrective refinement. Experiment 2 (**Fig.1b**) used the same task structure with only the high memory load condition, no resubmission or confidence procedures, and a response deadline manipulation: on deadline trials, participants were required to initiate their response within 3 seconds, with missed trials recorded as incorrect. This allowed us to test whether reward-related benefits persisted when response time was experimentally constrained, eliminating the possibility of SATs.

**Fig. 1.**
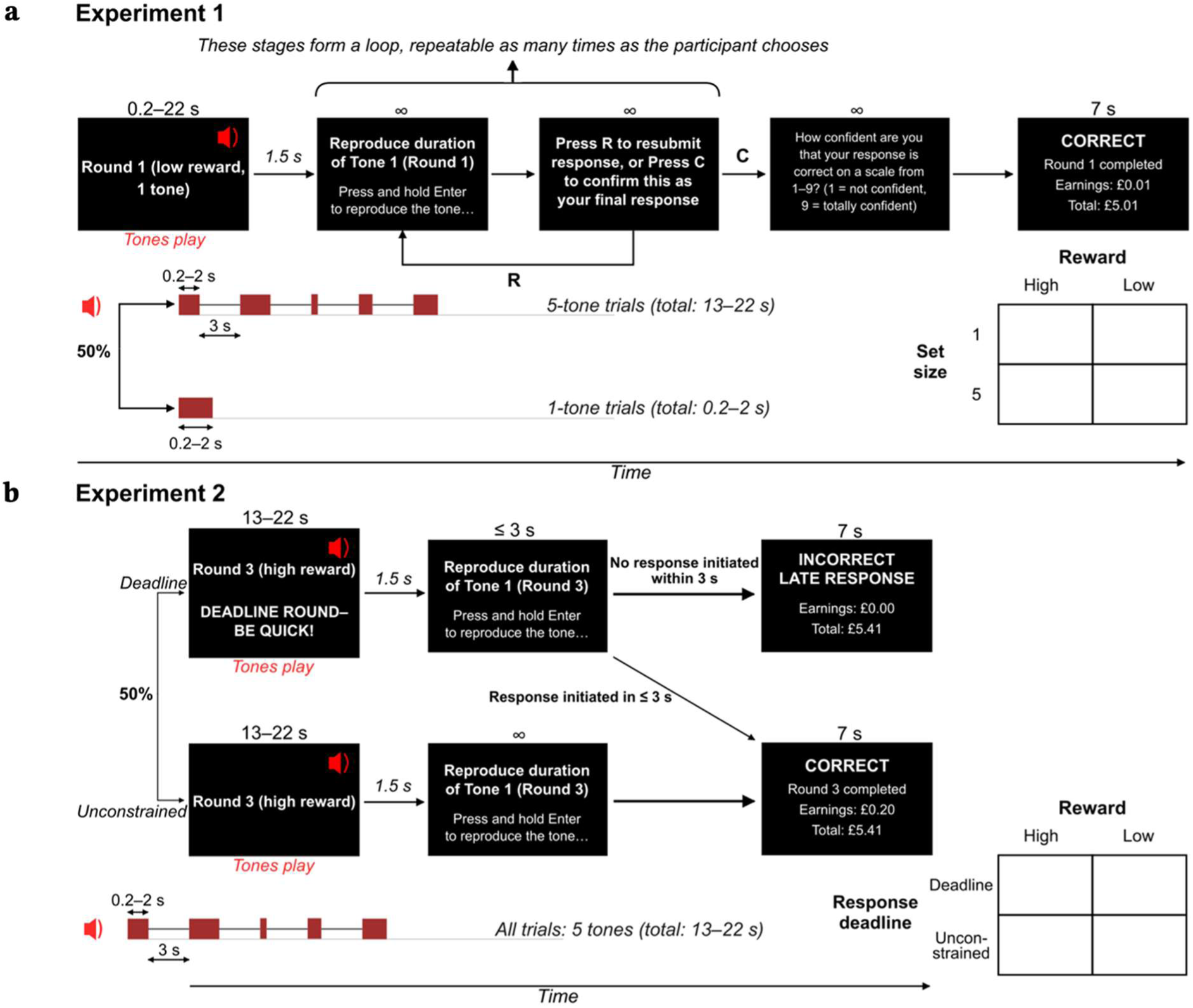
Auditory WM task structure across Experiments 1 and 2. (**a**) Experiment 1. Each trial began with an on-screen display indicating the trial number, memory load condition (one or five tones), and reward level (high, 20p; low, 1p), during which a tone sequence played (tones of 0.2–2.0 s each, separated by a 3-second silent interval in the five-tone condition). Fifty percent of trials were five-tone (high memory load) and 50% were one-tone (low memory load). After a 1.5-second retention interval, participants reproduced the duration of a probed tone by holding down the Enter key. Participants could then confirm (C) their response as final or resubmit (R) for another attempt, with the resubmission loop repeatable an unlimited number of times. After final confirmation, participants rated their confidence on a 1–9 scale and received feedback (7 s). The 2 × 2 factorial design manipulated reward and set size. (**b**) Experiment 2. Task structure was identical except that only the five-tone condition was used, resubmission and confidence procedures were omitted, and a response deadline manipulation was introduced. On 50% of trials, an on-screen alert (“DEADLINE ROUND-BE QUICK!”) accompanied the reward cue; on these trials, participants had to initiate their response within 3 seconds. If a response was initiated in time, it was submitted as in the unconstrained condition and CORRECT/INCORRECT feedback was displayed for 7 s; if no response was initiated within 3 seconds, no response was collected and the trial was recorded as incorrect with a LATE RESPONSE feedback screen. The remaining 50% of trials were unconstrained (no deadline), on which participants had unlimited time to respond. The 2 × 2 factorial design manipulated reward and response deadline.

### Experiment 1

#### Reward selectively improves WM performance under high memory load

If reward primarily influences response behaviour, improvements would be expected under both high and low memory load. In contrast, if reward enhances memory-based representations, improvements should be greatest under high load.

A 2 × 2 repeated-measures ANOVA revealed main effects of Reward (*M*_high_ = 0.156 s, *M*_low_ = 0.171 s), *F*(1,29) = 4.65, *P* = .040, ηp² = .14, and Memory Load (*M*_low_ = 0.144 s, *M*_high_ = 0.184 s), *F*(1,29) = 27.53, *P* = 1.3 × 10⁻⁵, ηp² = .49, and a significant Reward × Memory Load interaction, *F*(1,29) = 5.11, *P* = .032, ηp² = .15 (**Fig.2a**). Planned comparisons confirmed that, under high memory load, recall error was lower for high reward (*M* = 0.169 s) than low reward (*M* = 0.199 s), *t*(29) = −2.70, *P* = .011, whereas under low load, there was no difference between reward conditions (*M*_high_ = 0.143 s; *M*_low_ = 0.144 s), *t*(29) = −0.21, *P* = .836, BF₀₁ = 5.05.

**Fig. 2.**
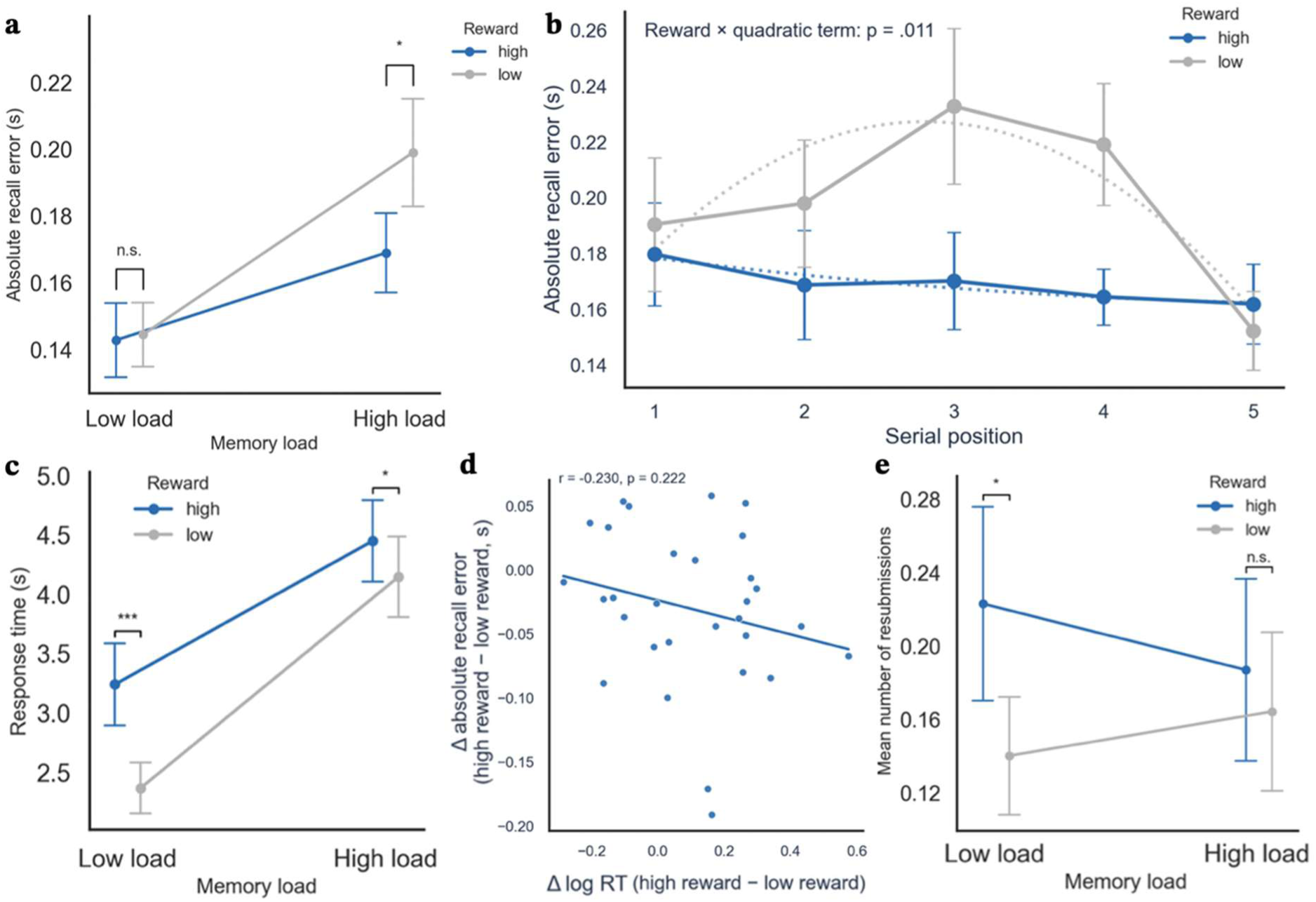
Reward-related improvements in Experiment 1 recall performance and their dissociation from response behaviour. (**a**) Recall error (s) as a function of reward condition (high, low) and memory load (low, high). Under low memory load, recall error did not differ between reward conditions; under high memory load, recall error was significantly lower under high reward compared with low reward. (**b**) Recall error (s) as a function of serial position, shown separately for high and low reward conditions in the high-load condition. Recall error was highest at middle serial positions, consistent with primacy and recency effects, and reward-related reductions in recall error were concentrated at these middle, high-interference positions and minimal at sequence edges. Dotted lines show the fitted quadratic mixed-effects model. (**c**) Response time (s) as a function of reward and memory load. Responses were slower under high reward, with the largest reward-related slowing under low memory load. (**d**) Individual-participant scatterplot of reward-related change in log response time (Δ log RT = high reward − low reward) against reward-related change in recall error (Δ error = high reward − low reward, s) under high memory load. No relationship was observed between reward-related changes in log response time and reward-related changes in recall error. Each point represents one participant, and the solid line shows the least-squares regression fit. (**e**) Mean number of resubmissions per trial as a function of reward and memory load. Reward increased the overall rate of resubmission, but this increase was significant only under low memory load. Error bars show standard errors of the mean. Asterisks indicate significant simple-effect comparisons (* *P* < .05, ** *P* < .01, *** *P* < .001; n.s., not significant).

Together, these results show that reward improves WM performance selectively under high memory load.

#### Reward selectively improves WM performance at serial positions with higher interference

In serial recall, items at the beginning and end of a sequence are typically remembered more accurately than middle items, due to primacy and recency effects^52–57^. If reward enhances memory-based representations, benefits should be greatest at middle positions with greater memory interference; if it primarily influences response behaviour, improvements should be uniform across positions.

Recall error benefits were greatest at middle positions (**Fig.2b**). A mixed-effects model revealed a significant quadratic effect of serial position, β = −0.007, *P* = .022, and a significant Reward × Quadratic interaction, β = 0.015, *P* = .011. Grouping serial positions into middle (2-4) and end (1 and 5) also confirmed a significant Reward × Position Type interaction, *F(*1,29) = 6.27, *P* = .018, ηp² = .18. Reward reduced recall error for middle positions, *t*(29) = 3.34, *P* = .002, but had no effect at ends, *t*(29) = 0.04, *P* = .971.

Together, these results show that reward flattens the serial-position curve by selectively improving WM performance at serial positions with greater memory interference.

#### Reward-related improvements are not explained by response behaviour

If reward improves WM performance by altering response behaviour rather than memory itself, the pattern of reward-related response changes should match the pattern of reward-related recall improvement, which was selective to high memory load. We examined the two response-behaviour markers, response slowing and resubmission rate, testing whether either paralleled the load-selective recall benefit.

Response times were log-transformed to correct for positive skew (see Methods). A 2 × 2 repeated-measures ANOVA revealed main effects of Reward (*M*_high_ = 3.826 s, *M*_low_ = 3.235 s), *F*(1,29) = 24.24, *P* = 3.14 × 10⁻⁵, ηp² = .46, and Memory Load (*M*_low_ = 2.792 s, *M*_high_ = 4.274 s), *F*(1,29) = 172.19, *P* = 1.00 × 10⁻¹³, ηp² = .86, and a significant Reward × Memory Load interaction, *F*(1,29) = 4.77, *P* = .037, ηp² = .14. Reward slowed responses under both loads, but planned comparisons revealed disproportionately greater slowing under low load (∼0.85 s slowing), *t*(29) = 4.64, *P* = .0001, BF₁₀ = 363.70, compared with high load (∼0.33 s slowing; **Fig.2c**), *t*(29) = 2.55, *P* = .016, BF₁₀ = 2.97; the largest reward-related slowing therefore occurred under the condition where no recall improvement was observed, dissociating from what a SAT account might predict (improvement with slowing down). Critically, when correlating per-participant reward-related changes in log RT and recall error under high load, individual differences in reward-related slowing did not predict individual differences in recall improvement, *r* = −0.230, *P* = .222, BF₀₁ = 2.16 (**Fig.2d**).

Resubmissions occurred on 13.2% (394/2,983) of trials and were used by 93.3% (28/30) of participants. A 2 × 2 repeated-measures ANOVA revealed a main effect of Reward (*M*_high_ = 0.204, *M*_low_ = 0.153), *F*(1,29) = 5.24, P = .030, ηp² = .15, but no effect of Memory Load, *F*(1,29) = 0.12, *P* = .737, BF₀₁ = 4.53, and no interaction, *F*(1,29) = 2.81, *P* = .104, BF₀₁ = 2.35 (**Fig.2e**). Planned comparisons further characterised this pattern; under low load, reward significantly increased resubmissions (*M*_high_ = 0.222, *M*_low_ = 0.141), *t*(29) = 2.75, *P* = .010, but no significant difference was observed under high load (*M*_high_ = 0.186, *M*_low_ = 0.166), *t*(29) = 0.80, *P* = .429, despite reward-related improvements in recall accuracy under high load. Critically, per-participant reward-related changes in resubmission rate did not predict per-participant changes in recall error under either load (high: *r* = −0.112, *P* = .557, BF₀₁ = 3.73; low: *r* = 0.014, *P* = .942, BF₀₁ = 4.39). Additionally, the Reward × Memory Load interaction on recall error was preserved when analysis was restricted to initial responses, before resubmission could contribute, *F*(1,29) = 6.71, *P* = .015, ηp² = .19 (Fig.S1).

Neither response-behaviour pattern shows the within-subject coupling with reward-related recall improvement that a response-criterion account would predict: response slowing and resubmission increases were both dissociated from recall benefits at the individual level, and the reward benefit was already present on initial responses. These convergent dissociations make response-behavioural accounts of the reward-related WM improvements unlikely.

#### Confidence ratings reveal metacognitively guided corrective refinement that dissociated from reward-driven memory enhancement

Confidence ratings were collected on every trial as a metacognitive index of response accuracy. If confidence merely reflects response-behavioural factors (response time or resubmission), the confidence-recall error relationship should disappear when these are controlled; if it reflects memory-based representation quality, this relationship should persist. Under the latter account, and assuming participants have metacognitive access to this internal memory signal, we may additionally expect that (i) low initial confidence predicts resubmission, (ii) resubmission improves both accuracy and confidence, and (iii) reward modulates confidence in parallel with its recall error effect, particularly under high memory load.

A linear mixed-effects model revealed that higher final confidence was associated with lower recall error (β = −0.026, *P* = 2.5 × 10⁻¹¹; **Fig.3a**). This relationship remained when controlling for response time (β = −0.024, *P* = 2.7 × 10⁻¹⁰), when controlling for whether responses were resubmitted (β = −0.023, *P* = 1.7 × 10⁻⁹), and when restricted to non-resubmitted trials (β = −0.023, *P* = 2.2 × 10⁻⁸). Confidence therefore tracked recall error independently of response time and resubmission, consistent with confidence indexing memory-based representation quality.

**Fig. 3.**
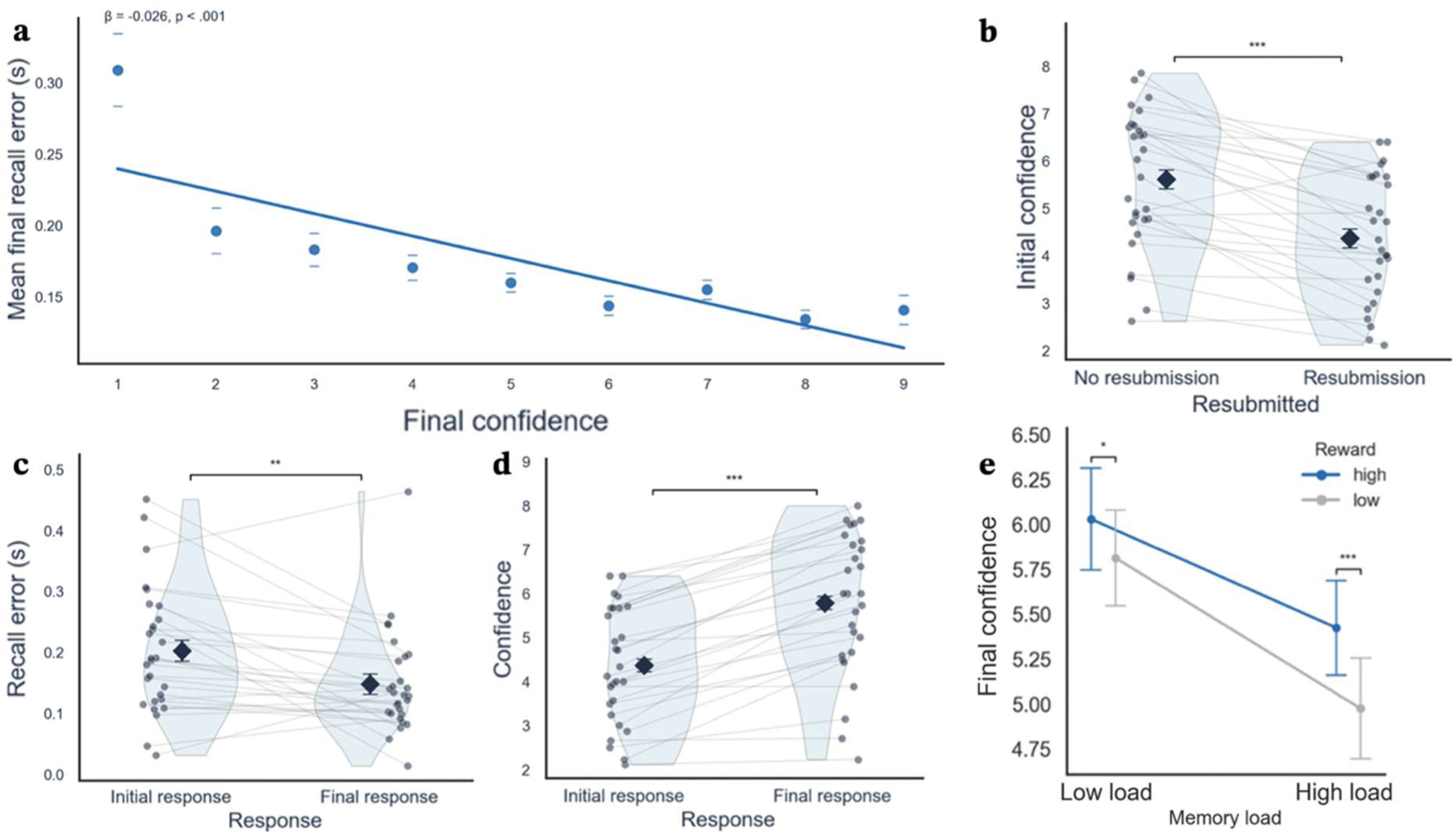
Relationships among confidence, recall error, and resubmission behaviour in Experiment 1. (**a**) Trial-level relationship between final confidence ratings (1-9 scale) and recall error (s). Higher confidence was associated with lower recall error. (**b**) Initial confidence on trials that were subsequently resubmitted versus not resubmitted. Trials that participants chose to resubmit were preceded by lower initial confidence, indicating that resubmissions were metacognitively triggered. (**c**) Initial and final recall error on trials with resubmission. Final recall error was significantly lower than initial recall error, indicating that resubmission reduced recall error. (**d**) Initial and final confidence ratings on trials with resubmission. Final confidence was significantly higher than initial confidence following resubmission. (**e**) Final confidence as a function of reward and memory load, showing higher confidence under high reward and a larger reward-related increase in confidence under high memory load. In panels b-d, grey dots and connecting lines show individual participants; dark diamonds and error bars show group means ± standard error of the mean. In panels a and e, error bars show standard errors of the mean. Asterisks indicate significant comparisons (* *P* < .05, ** *P* < .01, *** *P* < .001).

Lower initial confidence was associated with a higher rate of resubmission (β = −0.841, *P* = 2.8 × 10⁻⁴⁷; **Fig.3b**). On resubmitted trials, final recall error (*M* = 0.149 s) was lower than initial recall error (*M* = 0.204 s), *t*(27) = 3.25, *P* = .003 (**Fig.3c**), and final confidence (M = 5.79) was higher than initial confidence (*M* = 4.37), *t*(27) = −9.70, *P* = 2.7 × 10⁻¹⁰ (**Fig.3d**). Resubmissions were therefore triggered by low confidence and produced improvements in accuracy and subjective confidence.

Final confidence was also analysed as a function of reward and memory load (**Fig.3e**). A mixed-effects model revealed higher confidence under high than low reward (β = −0.218, *P*= .003), lower confidence under high than low memory load (β = −0.606, *P* = 5.3 × 10⁻¹⁷), and a significant Reward × Memory Load interaction (β = −0.216, *P* = .035), such that the reward-related increase in confidence was larger under high memory load. Simple-effect tests confirmed reward increased confidence at both low load (*t*(29) = 2.61, *P* = .014) and high load (*t*(29) = 4.19, *P* = 2.4 x 10⁻^4^); more so under high load.

Together, the observed pattern matched all four predictions: confidence tracked recall error independently of response behaviour, resubmissions were metacognitively triggered and outcome-improving, and reward increased confidence in parallel with recall error, most strongly under high memory load.

### Experiment 2

#### Reward-related improvements persisted under strict response deadlines

Experiment 1 suggested that response slowing is unlikely to explain motivational improvements; Experiment 2 tested whether improvements persist when decision time is constrained, using only the high memory load condition from Experiment 1 and introducing a response deadline (deadline, unconstrained).

Analysing log-transformed response times confirmed that the deadline manipulation effectively constrained response times (**Fig.4a**). A 2 × 2 repeated-measures ANOVA revealed a main effect of Deadline (*M*_deadline_ = 1.75 s, *M*_unconstrained_ = 2.47 s), *F*(1,29) = 26.34, *P* = 1.8 × 10⁻⁵, ηp² = .48, but no effect of Reward, *F*(1,29) = 0.0099, *P* = .922, BF₀₁ = 5.13, and no interaction, *F*(1,29) = 0.0564, *P* = .814, BF₀₁ = 5.00. Reward therefore did not slow responses under either condition. Response times under unconstrained conditions were shorter than in Experiment 1, indicating a global shift toward faster responding when deadline trials were present.

**Fig. 4.**
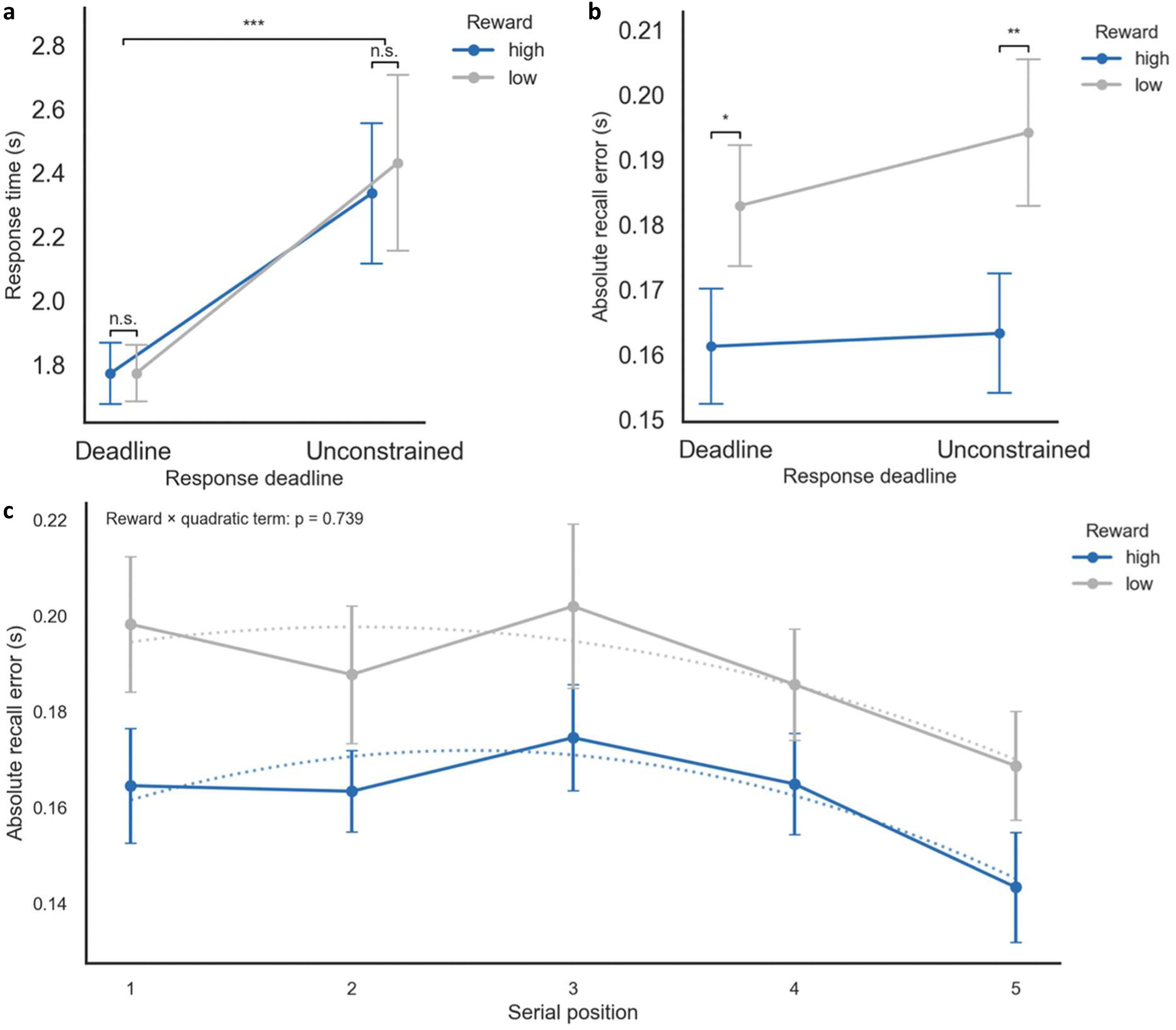
Reward, deadline, and serial position effects in Experiment 2. (**a**) Response time (s) as a function of reward and deadline. The deadline manipulation successfully constrained response time, and response times did not differ between reward conditions under either deadline. (**b**) Recall error (s) as a function of reward (high, low) and response deadline (fast, slow). Reward-related improvements in recall error were present under both deadline conditions and did not depend on the time available for responding. (**c**) Recall error as a function of reward and serial position (collapsed across deadline). Reward-related improvements were uniformly distributed across serial positions, in contrast to the interference-sensitive pattern observed in Experiment 1. Error bars show standard errors of the mean.

A 2 × 2 repeated-measures ANOVA on recall error revealed a significant main effect of Reward, *F*(1,29) = 16.24, *P* = .0004, ηp² = .36, no main effect of Deadline, *F*(1,29) = 0.95, *P* = .338, BF₀₁ = 3.19, and no interaction, *F*(1,29) = 0.45, *P* = .509, BF₀₁ = 4.07 (**Fig.4b**). Planned comparisons confirmed the reward benefit under both deadlines: recall error was lower high than low reward under strict deadlines (*M*_high_ = 0.162 s, *M*_low_ = 0.181 s), *t*(29) = −2.09, *P* = .046, and under unconstrained conditions (*M*_high_ = 0.163 s; *M*_low_ = 0.194 s), *t*(29) = −3.43, *P* = .0018. Reward therefore improved WM performance independently of response time, including under the globally compressed response regime imposed by the deadline.

Serial-position analyses were repeated to test whether the interference-sensitive pattern from Experiment 1 also manifested under temporal constraint. A 2 × 2 × 5 repeated-measures ANOVA with factors of reward, deadline, and serial position revealed significant main effects of Reward, *F*(1,29) = 16.31, *P* = .0004, ηp² = .36, and Serial Position, *F*(4,116) = 2.58, *P* = .041, ηp² = .08, but no Reward × Serial Position, *F*(4,116) = 0.10, *P* = .982, ηp² = .003 (all polynomial-contrast BF₀₁ > 4.6), Reward × Deadline, *F*(1,29) = 0.46, *P* = .503, ηp² = .02, BF₀₁ = 4.17, or three-way, *F*(4,116) = 0.26, *P* = .902, ηp² = .009, interaction, (all polynomial-contrast BF₀₁ > 4.2). A polynomial mixed-effects model confirmed significant linear (β = −0.0051, *P* = .018) and quadratic (β = −0.00375, *P* = .040) variation across positions, with a main effect of reward (β = −0.0237, *P* = .0127) that did not interact with these terms (all *P* ≥ .577 for polynomial x reward interaction terms). As shown in **Fig.4c**, reward-related improvements were uniformly expressed across serial positions under deadline constraints, unlike the interference-sensitive pattern observed in Experiment 1. This cross-experiment difference is considered further in the Discussion.

### Computational modelling suggests reward-related improvements in response precision

To characterise the mechanism underlying reward-related improvements, we fitted a Gaussian-uniform mixture model to signed recall errors (the signed residual from the participant-level regression; see Methods). This model decomposes responses into a memory-based component (Gaussian centred on the target duration, with standard deviation σ capturing spread of memory-based responses around the target) and a guessing component (uniform distribution across the response range, with mixture proportion, p_guess). Smaller σ indicates more precise memory-based responses, and the two parameters index distinct aspects of performance. If reward improves the quality of memory-based representations, this should manifest as reduced σ without changes in p_guess. Conversely, if improvements arise from changes in response behaviour, such as a lower rate of lapses or random responses, this should manifest as reduced p_guess.

Across both experiments, model comparison favoured the Gaussian-only model (Exp 1: ΔAIC = −0.72 to 1.00; ΔBIC = 0.49 to 2.21; Exp 2: ΔAIC = −0.01 to 1.35; ΔBIC = 1.21 to 2.57; Fig.S2), p_guess values were near zero for most participants (∼75-80%; Fig.S3a-b), and the mixture model was preferred in only a minority of cases (Fig.S3c-d). Representative fits captured empirical error distributions well (Fig.S4a-b). Behaviour was therefore well-described by a single memory-based process with minimal explicit guessing; the mixture model served as an explicit check on the guessing alternative.

#### Experiment 1

A 2 × 2 repeated-measures ANOVA on σ revealed significant main effects of Reward, *F*(1,29) = 6.31, *P* = .018, and Memory Load, *F*(1,29) = 23.32, *P* = 4.1 × 10⁻⁵, and a non-significant Reward × Memory Load interaction, *F*(1,29) = 3.03, *P* = .093, BF₀₁ = 1.35. As shown in **Fig.5a**, σ was lower under high than low reward, indicating improved memory-based precision, whereas σ was higher under high than low load (reduced memory-based precision under load). Although the interaction was not significant, the reward-related reduction in σ was numerically larger under high load. In contrast, neither reward, *F*(1,29) = 0.02, *P* = .898, BF₀₁ = 5.10, memory load, nor their interaction affected p_guess (**Fig.5b**; all *P* > .24, all BF₀₁ > 2.7).

**Fig. 5.**
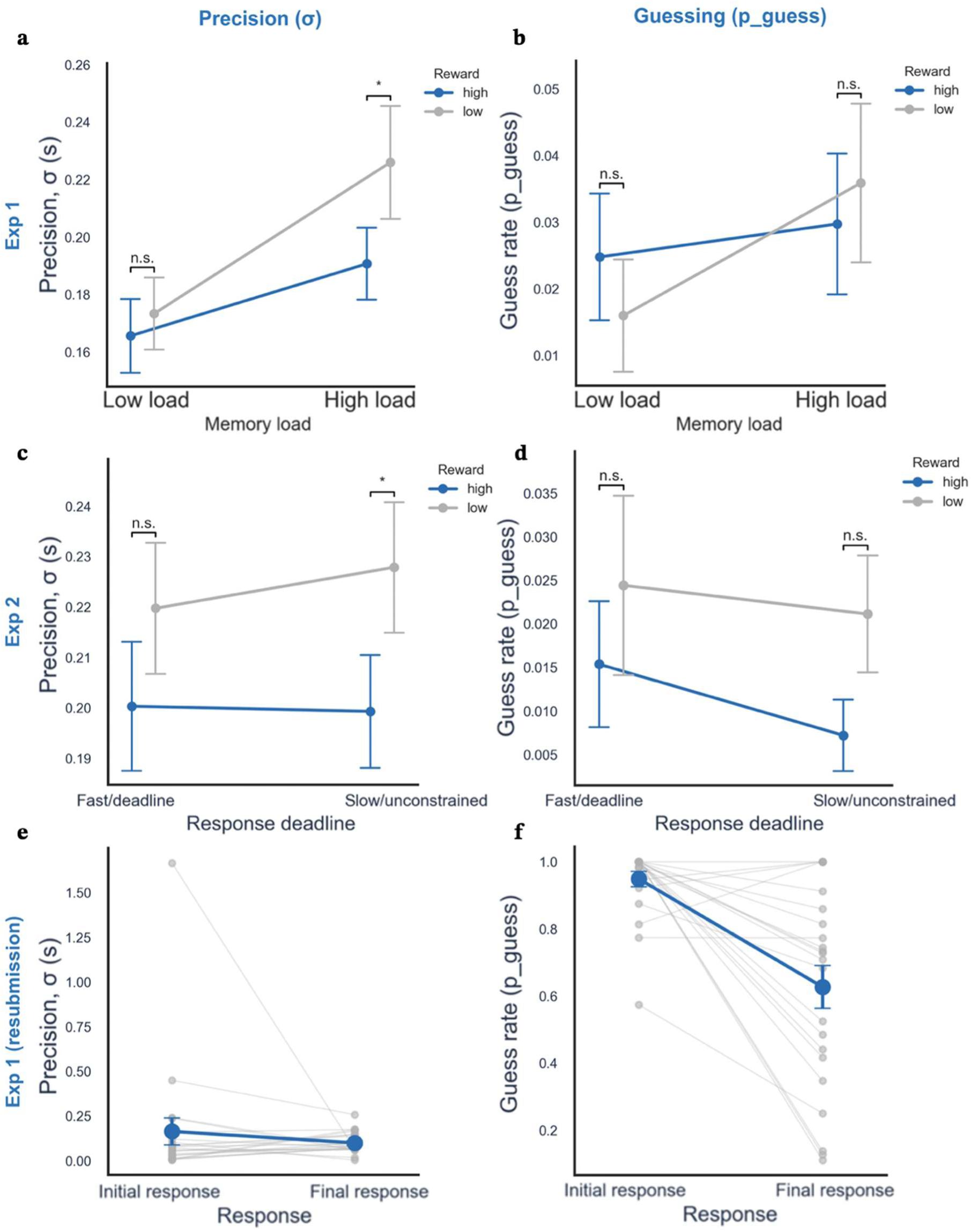
Mixture-model characterisation of reward and resubmission effects. Gaussian-uniform mixture model fitted to signed recall errors. Left column: precision parameter σ (units: seconds; smaller values indicate more precise memory-based responses). Right column: guessing parameter p_guess (probability of random guessing). (**a**) Experiment 1 σ as a function of reward and memory load; reward significantly reduced σ. (**b**) Experiment 1 p_guess as a function of reward and memory load; reward did not affect p_guess. (**c**) Experiment 2 σ as a function of reward and response deadline; reward again reduced σ, even under strict deadlines. (**d**) Experiment 2 p_guess as a function of reward and response deadline; reward did not affect p_guess. **(e, f)** Experiment 1 resubmission analysis: σ. (**e**) and p_guess (**f**) fitted separately to initial and final responses on trials with at least one resubmission (394 trials; *n* = 22 participants with usable initial and final fits). Resubmission did not reliably change σ (BF₀₁ = 3.26) but substantially reduced p_guess (from ∼0.95 to ∼0.63), indicating that resubmission converts initial guesses into memory-based responses rather than refining precision. In panels a–d, error bars show s.e.m. In panels e and f, grey lines and dots show individual participants; blue points show group means ± s.e.m. Asterisks indicate significant simple-effect comparisons (\**P* < .05, \*\**P* < .01, \*\*\**P* < .001; n.s., not significant).

#### Experiment 2

A 2 × 2 repeated-measures ANOVA on σ revealed a significant main effect of Reward, *F*(1,29) = 7.73, *P* = .009, no effect of Deadline, *F*(1,29) = 0.12, *P* = .729, BF₀₁ = 4.85, and no Reward × Deadline interaction, *F*(1,29) = 0.26, *P* = .614, BF₀₁ = 4.57. As shown in **Fig.5c**, σ was again lower under high than low reward, indicating that reward improved memory-based precision even under strict response deadlines, whereas the deadline itself did not affect σ. Neither reward, *F*(1,29) = 2.29, *P* = .141, BF₀₁ = 1.85, deadline, nor their interaction affected p_guess (**Fig.5d**; all *P* > .48, BF₀₁ > 4.0).

#### Resubmission reduced guessing rather than improving precision (Experiment 1)

Because resubmissions were permitted only in Experiment 1, we further asked what changes about a response when participants choose to resubmit. Of the 13.2% of trials (394 of 2,983; 28 of 30 participants) with at least one resubmission, we fitted the mixture model separately to initial and final responses, comparing σ and p_guess within participants. Collapsing across reward and memory load, resubmission did not reliably change σ (*M*_initial_ = 0.164 s, *M*_final_ = 0.099 s), *t*(21) = 0.85, *P* = .408, BF₀₁ = 3.26 (**Fig.5e**), but substantially reduced p_guess (*M*_initial_ = 0.95, *M*_final_ = 0.63), *t*(21) = 4.75, *P* = 1.1 × 10⁻⁴, BF₁₀ = 252 (**Fig.5f**). The p_guess reduction held across all subgroups (low reward BF₁₀ = 29.5; high reward BF₁₀ = 1.68; low load BF₁₀ = 4.58; high load BF₁₀ = 4.73), whereas the σ reduction was not reliable in any subgroup (all *P* ≥ .11, all BF₀₁ ≥ 1.21; anecdotal reduction only under high reward, BF₁₀ = 1.93). Resubmission therefore reduces p_guess but does not reliably reduce σ, dissociating from the reward-related pattern.

#### Robustness of the reward effect on improving precision

The reward effect on σ remained consistent after excluding extreme responses (top and bottom 1% of trials; Fig.S5a-b) and when varying parameter bounds (Fig.S5c-d). To rule out an artefact of differential resubmission rates, we refitted the main analysis to non-resubmitted trials only (86.8% of Experiment 1 trials); the reward effect on σ was slightly stronger, *F*(1,29) = 7.70, *P* = .010, BF₁₀ = 4.68 (Fig.S6a), while p_guess remained unaffected, *F*(1,29) = 1.03, *P* = .319, BF₀₁ = 3.22 (Fig.S6b), confirming the reward-related precision benefit is not an artefact of resubmission behaviour.

#### Summary of modelling results

Reward therefore reduces σ without changing p_guess, indicating that reward-related improvements reflect more precise memory-based responses rather than reduced guessing. The resubmission analysis shows that this precision benefit cannot be attributed to resubmission: resubmission itself reduced p_guess (converting initially poor responses into memory-based responses) without reliably changing σ, dissociating from the reward-related improvements. This dissociation also confirms that σ and p_guess are separable in this dataset, so reward’s null effect on p_guess is a genuine null rather than a limitation of the measure. Together, these findings support a memory-representation account of reward-related WM improvements.

## Discussion

Across two experiments and converging computational modelling, the present findings indicate that motivational improvements in equal-value WM items are better explained by enhanced memory-based representations than by a change in response criterion, i.e. by speed-accuracy trade-offs or other changes in response behaviour. In Experiment 1, although reward slowed responses at both memory loads, slowing occurred disproportionately more under low load, where no recall benefit occurred, than under high load, where recall benefits were concentrated. Similarly, individual differences in reward-related slowing did not predict individual differences in reward-related recall improvement. Both patterns are difficult to reconcile with a speed-accuracy trade-off account that predicts improvement with slowing down. In Experiment 2, motivational benefits persisted under strict response deadlines and occurred without detectable reward-related slowing despite substantial overall reductions in response time. Together, these dissociations make it unlikely that increased response caution and deliberation^39,42,43,58^, could explain the observed pattern.

Instead, motivational incentives appear to improve WM performance by enhancing the quality of memory-based representations guiding responses. In Experiment 1, motivational benefits emerged selectively under high memory load, being disproportionately expressed at middle serial positions typically associated with greater retrieval competition^45,52,59,60^. Under low load, where WM demand is minimal, no reward-related improvement was observed (**Fig.2a**), consistent with motivational benefits depending on the memory component of the task rather than response behaviour changes. Confidence analyses further supported this interpretation. Trial-level confidence tracked recall error *independently* of response time and resubmission, consistent with confidence indexing the quality of memory-based representations, rather than response time or post-response adjustment. Low initial confidence prompted resubmissions that reduced recall error and increased confidence, indicating a metacognitively-guided refinement process in which participants adjust responses toward an underlying memory-based representation. Critically, this refinement did not itself drive the reward-related recall benefit: reward increased resubmissions primarily under *low* load, where recall did not improve, and did not increase them under high memory load, where recall benefits were concentrated. Corrective refinement may therefore reflect adjustments toward an already motivationally-improved memory representation, rather than producing the improvement. Consistent with this, reward-related increases in confidence were larger under high load (**Fig.3e**), paralleling the recall benefit (**Fig.4b**). In Experiment 2, motivational benefits persisted under strict response deadlines, indicating that benefits do not require additional decision time. Together, these convergent lines of evidence suggest that motivation can indeed enhance memory-based representations, rather than simply altering response behaviour.

Extending this interpretation, mixture-model analyses revealed a dissociation at the parameter level: reward increased precision, without changing the probability of guessing, whereas on resubmitted trials, resubmission substantially reduced guessing rates without reliably improving precision (**Fig.5**). This dissociation maps directly onto the metacognitive account outlined above. Reward improves the internal memory signal itself, sharpening the memory-based response distribution without altering the rate at which memory is accessed. Resubmission, however, operates at the level of response commitment: participants use metacognitive access to the signal to detect low-quality initial responses and reallocate to further attempts, converting initially poor responses into memory-based responses (reduced probability of guessing) without further refinement of the retrieved memory (unchanged precision). Reward and resubmission therefore act at different stages of the retrieval-response process: reward improves the quality of the underlying representation, whereas resubmission changes whether that representation is used at retrieval. The opposite direction of the two effects is important: it establishes precision and guessing as dissociable within the current dataset, and reward’s null effect on guessing is informative, rather than a limitation of the measure.

Notably, the serial-position pattern of motivational improvements differed between the two experiments. In Experiment 1, benefits were concentrated at middle serial positions associated with greater retrieval competition (**Fig.2b**); in Experiment 2, where retrieval was temporally constrained, enhancement was expressed more uniformly across positions (**Fig.4c**). This difference is consistent with a broader class of accumulation-model accounts of retrieval, in which interference-sensitive expression of representational benefits depends on retrieval being able to engage in competitive selection between memory traces; this is characterised as a time-dependent process, where a differential advantage of higher-quality representations over weaker ones accrues progressively as retrieval dynamics resolve^4,61,62^. Under flexible retrieval, this differential advantage can fully resolve, concentrating benefits at high-competition items. Under temporal constraints, competing traces may be selected before differential read-out fully resolves^58^, so that representational benefits are expressed more uniformly across positions. Consistent with this, response times in Experiment 2 were shorter even on nominally unconstrained trials than in Experiment 1, indicating a globally compressed retrieval regime^44^. Critically, both patterns are consistent with changes in the quality of memory-based representations rather than changes in response behaviour.

While the present findings constrain the mechanism by which motivational improvements in WM occur, they do not isolate the stage within the WM process at which this enhancement arises. Since reward was cued before stimulus presentation, motivational state could influence encoding^18^, maintenance, or retrieval^63,64^, and stage-specific manipulations will be needed to distinguish these. Beyond the stage-level question, several candidate mechanisms could underlie motivational enhancement of memory-based representations.

One possibility is that motivational incentives enhance memory-based representation quality by increasing their signal-to-noise ratio^65–67^. Computationally, this could involve either suppressing background noise or amplifying task-relevant representations through increased neural gain^68–70^ or attentional amplification^71–73^. At the circuit level, such modulation may reflect dopaminergic stabilisation of prefrontal attractor states^74,75^, resulting in increased signal-to-noise ratio^74,76^ and more distinct representations of competing items. This is consistent with biased-competition accounts of WM^71,77^, and the present interference-sensitive pattern of reward-related improvement.

An alternative directly supported by our data, is that motivational incentives reduce variability in memory-based representations rather than increasing their mean strength. Resource models emphasise trial-to-trial variability in representational precision as a key performance constraint^5,23,25^, and our data showed that reward selectively reduced the variability of memory-based responses (σ) without affecting guess rate. This stabilisation could arise through more consistent encoding or maintenance dynamics, decreased lapse probability^27,78^, reduced susceptibility to interference^59^, or enhanced preparatory control or attentional engagement^12,79^.

Motivational incentives may also act at a higher level by modulating adaptive control allocation. Resource-rational^80,81^ and effort-allocation accounts propose that motivational incentives increase the investment of cognitive resources when expected benefits outweigh costs^12,28^. Reward cues available before stimulus presentation may alter preparatory control states^14^, consistent with frameworks in which motivation promotes proactive control^82,83^. In this view, incentives do not introduce a distinct computational mechanism, but rather increase the consistency and effectiveness with which existing processes are deployed^84^, potentially by enhancing encoding fidelity, stabilising maintenance, or improving retrieval selection. Motivational incentives may also promote more structured encoding strategies, such as improved rehearsal organisation or chunking of sequential material^1^, reflecting a strategic expression of increased control investment.

These candidate mechanisms are not mutually exclusive. Motivational incentives may simultaneously increase neural gain, reduce representational variability, and alter control allocation. While the present data constrain the computational outcome to enhancement of memory-based representations, distinguishing between signal amplification, variability reduction, and adaptive control mechanisms will require neurophysiological or pharmacological approaches targeting dopaminergic systems^75^, or hierarchical modelling that explicitly characterise trial-level variability^3,5,27^.

The present findings bear on broader theories of motivated cognition and cognitive control. Contemporary resource-rational and expected-value-of-control frameworks^12,28,80^ propose that motivational incentives regulate cognitive performance according to expected value, typically through adjustments in control allocation or effort investment^14^, allowing for improvements in representational precision^5,25^ when anticipated benefits justify the costs of enhanced processing; value signals have been shown to influence internal representations early in processing^85^, consistent with motivational effects not requiring prolonged deliberation. Within WM literature, motivational enhancements have often been attributed to selective prioritisation of certain items^15,16,18^ or, in equal-value contexts where prioritisation is not possible, to changes in response behaviour during retrieval^39^. The present findings refine this interpretation: by demonstrating that motivational benefits are better explained by enhancing memory-based representations, rather than changes in response behaviour, we argue that value-based modulation operates at the level of the representations themselves. Importantly, because all items were equally valued, this enhancement is unlikely to reflect redistribution of limited resources across items. The present findings therefore extend contemporary motivated-control theory by providing behavioural and computational evidence that value signals influence the fidelity of internal representations, not merely the policies governing their deployment. Why, then, do our results differ from earlier studies that found no equal-value reward benefit^38^ or attributed benefits entirely to response behaviour^39^?

Several methodological differences are likely relevant. Grogan et al.^39^ did not manipulate memory load within experiments, whereas our data show that reward benefits emerge selectively at higher load (**Fig. 2a**); studies without an explicit load manipulation may miss the load-dependence of the effect entirely. Second, two of van den Berg et al.’s^38^ three experiments used a between-subjects reward manipulation, which is likely to under-estimate motivational effects because reward valuation is inherently relative. Participants who never experience the alternative reward level lack a within-subject contrast against which to calibrate effort. Finally, both prior studies used visual orientation reproduction, whereas we used auditory duration reproduction; the extent to which our findings generalise across sensory modalities warrants further investigation.

Some limitations warrant consideration. First, because Experiment 2 included only high memory load, we could not directly test whether the load-specificity of reward effects observed in Experiment 1 is preserved under temporal constraint. The absence of any reward effect under low load in Experiment 1 makes it unlikely that one would emerge under low load with a deadline, but the three-way Reward × Load × Deadline interaction was not directly tested. Second, the studies were not pre-registered; however, sample sizes were determined a priori (see Methods), sensitivity analyses confirmed 80% power to detect medium-sized within-subjects effects (Cohen’s *d* = 0.53 at α = .05), and Bayes Factors reported alongside null effects quantify their evidential strength. Third, the deadline manipulation in Experiment 2 altered overall response dynamics, with response times shorter even in nominally unconstrained trials than in Experiment 1. Although this contextual shift was theoretically informative, future work orthogonally manipulating deadline pressure while holding task context constant would clarify how time constraints interact with memory-based representations. Finally, the present conclusions rely on behavioural and computational inference rather than direct neural measurement; neurophysiological or pharmacological approaches are required to distinguish between increased neural gain, reduced representational variability, dopaminergic modulation of cortical stability, or related processes.

In summary, we addressed a central question in motivated cognition: whether motivational improvements in equal-value working memory reflect changes in response behaviour or improvements in the quality of memory-based representations. Our findings support the latter. Reward-related improvements persisted under strict response deadlines and did not track the pattern of reward-related slowing, indicating that SATs are unlikely to account for the observed benefits. Instead, motivational incentives appear to directly enhance the fidelity and stability of information maintained in working memory.

## Methods

The study consisted of two experiments examining the effects of motivational incentives on working memory (WM) performance under equal-value conditions.

### Experiment 1

#### Design

Experiment 1 used a 2 × 2 within-subjects design manipulating reward (high vs low) and memory load (high vs low). The reward manipulation varied the amount of money participants could earn for accurate responses (20p vs 1p). The memory load manipulation varied the number of auditory items participants were required to maintain in WM (five tones vs one tone). The primary outcome measure was recall error.

#### Participants

Sample size (*n* = 30) was matched to prior comparable WM studies using within-subjects reward manipulations^39^, for which this sample size was determined to be sufficient to detect medium effects at conventional significance levels. A sensitivity analysis confirmed that *n* = 30 provided 80% power to detect within-subjects effects of Cohen’s *d* = 0.53 at α = .05 (two-tailed) for paired-samples comparisons, consistent with Grogan et al.’s^39^ prior justification. 30 participants (22 male, 8 female; aged 18-60 years) were recruited. All participants provided written informed consent prior to participation. The study was ethically approved by the University of Oxford Medical Sciences Interdivisional Research Ethics Committee and was performed in accordance with the Declaration of Helsinki and all relevant institutional guidelines and regulations. Participants received a base payment of £5.00, with additional performance-based monetary rewards.

#### Stimuli

The experiment consisted of 100 trials in a fully crossed 2 × 2 design (reward × memory load), with 25 trials per condition, presented in a random order. In the high memory load condition, participants heard a sequence of five tones; in the low memory load condition, a single tone was presented. Reward level was manipulated independently, with trials designated as either high reward or low reward. Each tone consisted of a 2000 Hz sine wave with a randomly generated duration between 0.2 and 2.0 seconds. In the five-tone (high memory load) condition, consecutive tones were separated by a 3-second silent interval; no such interval applied in the one-tone (low memory load) condition.

#### Apparatus

Computer events were driven by a 2020 Apple MacBook Pro running macOS Monterey (version 12.7.1). The experiment was implemented in Python (version 3.11) and executed using PyCharm. All statistical analyses were conducted using Python with standard scientific libraries (NumPy, SciPy, statsmodels). Participants completed the experiment in a quiet room, with the experimenter present face-to-face.

#### Procedure

Each trial began with an on-screen display indicating the trial number, memory load condition, and reward level (**Fig.1a**). Participants then heard either a single tone (low load) or a sequence of five tones (high load). After a 1.5-second retention interval, participants were prompted to reproduce the duration of a specified tone from the sequence. Participants reproduced the target duration by holding down the Enter key for the perceived duration of the target tone. The length of the keypress constituted the response. Following this initial response, participants could either confirm their response or choose to resubmit it. If resubmission was selected, participants repeated the reproduction process. This option could be used an unlimited number of times, allowing iterative refinement prior to final confirmation.

After confirming their response, participants provided confidence ratings. On trials with a single response, participants reported confidence in their response accuracy on a scale from 1 (not confident at all) to 9 (totally confident). On trials involving resubmissions, participants provided separate confidence ratings for their initial and final responses.

Following confidence ratings, participants received trial feedback indicating whether their final response fell within a predefined accuracy window (correct/incorrect), the reward earned on that trial, and cumulative earnings. Feedback was displayed for 7 seconds before the next trial began. A response was considered correct if the absolute difference between the response and target duration satisfied |Response − A| ≤ (0.15 + 0.1A), where A denotes the target duration. The proportional term (0.1A) accommodates Weber-law scaling of temporal precision, whereby just-noticeable differences scale with stimulus duration^86,87^. This threshold was set a priori and verified in the first five participants (62.2% of trials correct across 500 trials), after which data collection continued unchanged. Only final responses were used to determine accuracy.

Prior to the main experiment, participants completed four practice trials (one per condition) to familiarise themselves with the task. Practice trials did not contribute to earnings. The full experiment lasted approximately 50 minutes.

#### Behavioural Measures

**<u>Recall Error:</u>** The primary measure of WM performance was error in reproduced duration. Previous studies using auditory tones as stimuli have shown a systematic bias whereby shorter tones are overestimated, and longer tones are underestimated^52,86,87^, consistent with Vierordt’s law^88^. To account for this bias, a linear regression was fit to the relationship between target and response durations for each participant (Fig.S7). Recall error was defined as the absolute residual from this regression, providing a measure of deviation from the expected response independent of central tendency bias.

**<u>Response Time:</u>** Response time was defined as the interval between presentation of the response prompt and the onset of the participant’s keypress initiating their final response. Inspection of the response time distribution revealed extreme outliers corresponding to trials exceeding 30 seconds, likely reflecting lapses in attention or disengagement from the task. Across all participants, 17 such trials (0.567% of trials) were excluded.

**<u>Number of Resubmissions:</u>** Participants were allowed to resubmit their responses before final submission; the number of resubmissions per trial was recorded, serving as an index of corrective refinement.

**<u>Confidence ratings:</u>** Confidence ratings were used as a trial-by-trial subjective measure of response certainty. Ratings were provided on a scale from 1 (not confident at all) to 9 (totally confident).

**<u>Signed recall error:</u>** Signed recall error was defined as the difference between the participant’s reproduced duration and the target duration (based on the same linear regression per participant as for recall error), with negative values indicating underestimation and positive values indicating overestimation. This measure was used for mixture modelling.

#### Statistical Analysis

All statistical analyses were conducted in Python using standard scientific libraries. Assumptions of parametric tests were assessed via Shapiro-Wilk tests on participant-level cell means and mixed-model residuals. Response times exhibited positive skew and were log-transformed prior to analysis, which restored normality of the residuals. Recall error was by definition positive because it was defined as an absolute residual, and therefore followed a half-normal-like distribution by construction; consistent with this, Shapiro-Wilk tests identified some cells as non-normal despite the underlying signed error distributions being well-behaved. Because repeated-measures ANOVA is robust to moderate normality violations at *n* = 30, and because parallel analyses on log-transformed recall error (which restored normality across all cells, Shapiro-Wilk *W* > 0.94, all *P* > .07) yielded a consistent pattern of results, untransformed recall error is reported throughout for interpretability. Bayes Factors are reported alongside all key null and near-threshold effects, providing inference that does not rely on strict distributional assumptions. The number of resubmissions was analysed at the participant level as mean values per condition.

Effects of reward and memory load on recall error, RT, and resubmission data were analysed using two-way repeated-measures ANOVAs. Where appropriate, planned pairwise comparisons were conducted using paired-sample *t*-tests. Associations between reward-related changes in response time and recall error were assessed using Pearson correlation coefficients. Reward-related change scores (Δ) were computed as high reward minus low reward for each measure. Bayes Factors were computed using pingouin^89^, with a default Jeffreys-Zellner-Siow prior^90^. For interactions in the 2 × 2 × 5 ANOVA, we computed Bayes Factors on participant-level contrasts (difference-of-differences for 2-way interactions; linear and quadratic polynomial contrasts of the reward effect across positions for interactions involving serial position). Values of BF₀₁ > 3 are conventionally interpreted as moderate evidence for the null.

Because serial position varied only in high-load trials, recall error across serial positions was analysed using linear mixed-effects models restricted to only these trials. Models included linear and quadratic polynomial predictors of serial position, fixed effects of reward and serial-position predictors, and participant-level random intercepts. Additional analyses grouped serial positions into middle positions (positions 2-4) and end (positions 1 and 5) categories, which were analysed using repeated-measures ANOVA with factors of reward (high, low) and position type (middle, end).

Confidence ratings were analysed using linear mixed-effects models predicting recall error from standardised trial-level confidence ratings. Response time was also separately included as a covariate. Logistic regression models were used to predict the probability of resubmission from initial confidence ratings. Paired-sample t-tests compared initial and final recall error, as well as initial and final confidence ratings, on trials involving resubmissions.

All statistical tests were two-tailed with α = .05.

### Experiment 2

#### Design

Experiment 2 used a 2×2 within-subjects design manipulating reward (high vs low) and response deadline (deadline vs unconstrained). The reward manipulation was identical to Experiment 1 (20p vs 1p). The deadline condition varied the amount of time participants had to initiate their response (3-second deadline vs unconstrained). The primary dependent variable was recall error. Because the reward effect on recall error in Experiment 1 was selective to high memory load, Experiment 2 used only the high load condition (sequences of five tones) to maximise statistical power for testing the central question of whether reward-related improvements persist under deadline constraints.

#### Participants

30 participants (19 male, 11 female; aged 18-60 years) were recruited, of whom 21 had also participated in Experiment 1. Sample-size rationale, sensitivity considerations, ethical approval, and compensation were as described for Experiment 1.

#### Stimuli

The experiment consisted of 100 trials in a fully crossed 2 × 2 design (reward × deadline), with 25 trials per condition, presented in a randomised order. All trials involved sequences of five tones. Each tone was a 2000 Hz sine wave with a duration randomly sampled between 0.2 and 2.0 seconds. Consecutive tones were separated by a 3-second silent interval, as in Experiment 1.

#### Apparatus

Apparatus was identical to that used in Experiment 1.

#### Procedure

The trial structure was similar to that of Experiment 1, with the following modifications. All trials involved sequences of five tones, and the resubmission and confidence-rating procedures were omitted. A response deadline manipulation was introduced.

Each trial began with an on-screen display indicating the trial number, reward level, and if relevant, an alert to a deadline being present. Participants then heard a sequence of five tones. After a 1.5-second retention interval, participants were prompted to reproduce the duration of a specified tone from the sequence. Participants reproduced the target duration by holding down the Enter key for the perceived duration of the tone. The length of the keypress constituted the response.

In the unconstrained condition, responses were automatically submitted upon key release. In the deadline condition, participants were required to initiate their response within 3 seconds. If the deadline was met, responses were submitted as in the unconstrained condition. If the deadline was not met, the trial was recorded as incorrect and no response was collected (**Fig.1b**). Feedback was then provided, indicating response accuracy (correct/incorrect), whether the response was too slow (if applicable), the reward earned on that trial, and cumulative earnings. Feedback was displayed for 7 seconds before the next trial began.

Trials in which no response was initiated within the deadline were excluded from analysis. Across all participants, 57 trials (1.9%) were excluded on this basis.

Prior to the main experiment, participants completed four practice trials (one per condition) to familiarise themselves with the task. Practice trials did not contribute to earnings.

#### Behavioural measures

**<u>Recall error:</u>** See experiment 1.

**<u>Response time:</u>** Response time was defined as the interval between presentation of the response prompt and the onset of the participant’s keypress initiating the response. Inspection of the response time distribution across trials revealed extreme outliers corresponding to trials exceeding 20 seconds, likely reflecting lapses in attention or disengagement from the task. Therefore, across all participants, 5 trials (0.167% of trials) were excluded from analysis due to response times exceeding 20 seconds.

**<u>Signed recall error:</u>** See experiment 1

#### Statistical analysis

Conducted as in Experiment 1 for recall error, response time, and serial position analysis.

### Mixture modelling of recall errors

Signed recall errors were analysed using a Gaussian-uniform mixture model. This model decomposes responses into two components: a memory-based component reflecting noisy recall of the target, and a guessing component reflecting random responding.

Errors were modelled as arising from a mixture of (i) a Gaussian distribution centred on zero, representing responses based on noisy memory of the target, and (ii) a uniform distribution spanning the full range of possible errors, representing random guesses. The probability density of an observed error, *x*, was therefore given by:

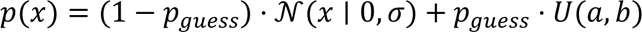

where *σ* represents the standard deviation of the Gaussian distribution (capturing memory precision), *p*_guess_represents the probability of random guessing, and *U*(*a*, *b*) denotes a uniform distribution across the range of possible recall errors. Lower values of *σ* correspond to more precise memory representations.

The model was fit separately for each participant and condition using maximum likelihood estimation. Each participant contributed approximately 25 trials per condition, yielding a total of 2,983 trials in the adjusting experiment and 2,943 trials in the deadline experiment.

Parameter optimisation was performed using bounded numerical optimisation (L-BFGS-B), with constraints σ > 0 and 0 ≤ p_guess ≤ 1. To ensure numerical stability, likelihood values were bounded away from zero.

Model fit was compared to a Gaussian-only model (i.e. without a guessing component) using the Akaike Information Criterion (AIC) and Bayesian Information Criterion (BIC), computed separately for each participant and condition.

To assess robustness, analyses were repeated under alternative assumptions regarding the bounds of the uniform distribution (±10-20% of the observed error range) and after excluding extreme responses (top and bottom 1% of signed errors).

Parameter estimates (*σ* and *p*_guess_) were analysed using repeated-measures ANOVAs with factors corresponding to the experimental manipulations in each experiment (reward × memory load in Experiment 1; reward × deadline in Experiment 2). All statistical tests were conducted at the participant level.

## Transparency note

The studies reported here were not pre-registered. Sample sizes were determined a priori based on prior comparable studies using within-subjects reward manipulations in continuous-reproduction WM tasks (see Participants subsections), and sensitivity analyses were performed to confirm that this sample size provided adequate statistical power (80% power to detect within-subjects effects of Cohen’s *d* = 0.53 at α = .05 for paired-samples comparisons). All measures, manipulations, and data exclusions are reported in the main text.

## Data availability

The anonymised data and analysis code supporting the findings of this study will be made publicly available on the Open Science Framework (https://osf.io/) upon publication of the manuscript. In the interim, data are available from the corresponding author upon request.

## Acknowledgements

The authors thank the Department of Experimental Psychology, University of Oxford for supporting the work. SGM was funded by an MRC project grant UKRI2552 and by the National Institute for Health and Care Research (NIHR) Oxford Health Biomedical Research Centre (BRC).

## Author contributions

V.N. conceived the study, designed and conducted the experiments, analysed the data, and wrote the manuscript. O.A. contributed to aspects of the analytical methodology and provided associated analysis code. S.M. and E.M. supervised the project, provided input on study design and interpretation, and reviewed and edited the manuscript. All authors approved the final version of the manuscript.

