## Supplementary material for "Motivational incentives improve working memory performance by enhancing memory-based representations rather than altering response strategies"

Supplementary Information

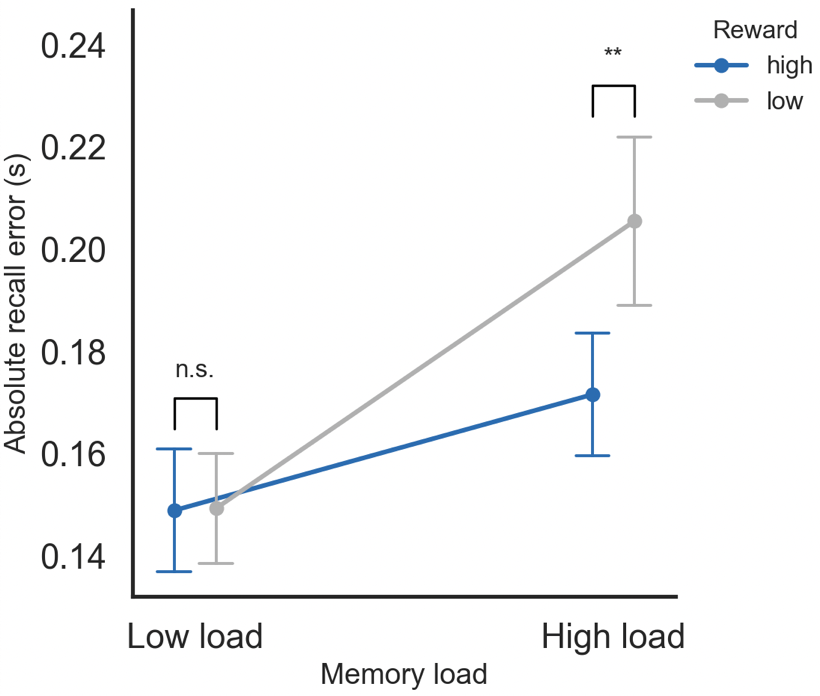

**Fig.S1. Reward × Memory Load effect on absolute recall error restricted to initial responses (Experiment 1).** Absolute recall error (s) computed using only participants' initial responses, before any resubmission could contribute, as a function of reward (high, low) and memory load (high, low). The Reward × Memory Load interaction was significant, F(1,29) = 6.71, P = .015, ηp² = .19, matching the pattern observed for final recall error, indicating that reward-related improvements were present prior to any refinement through resubmission. Error bars show standard errors of the mean.

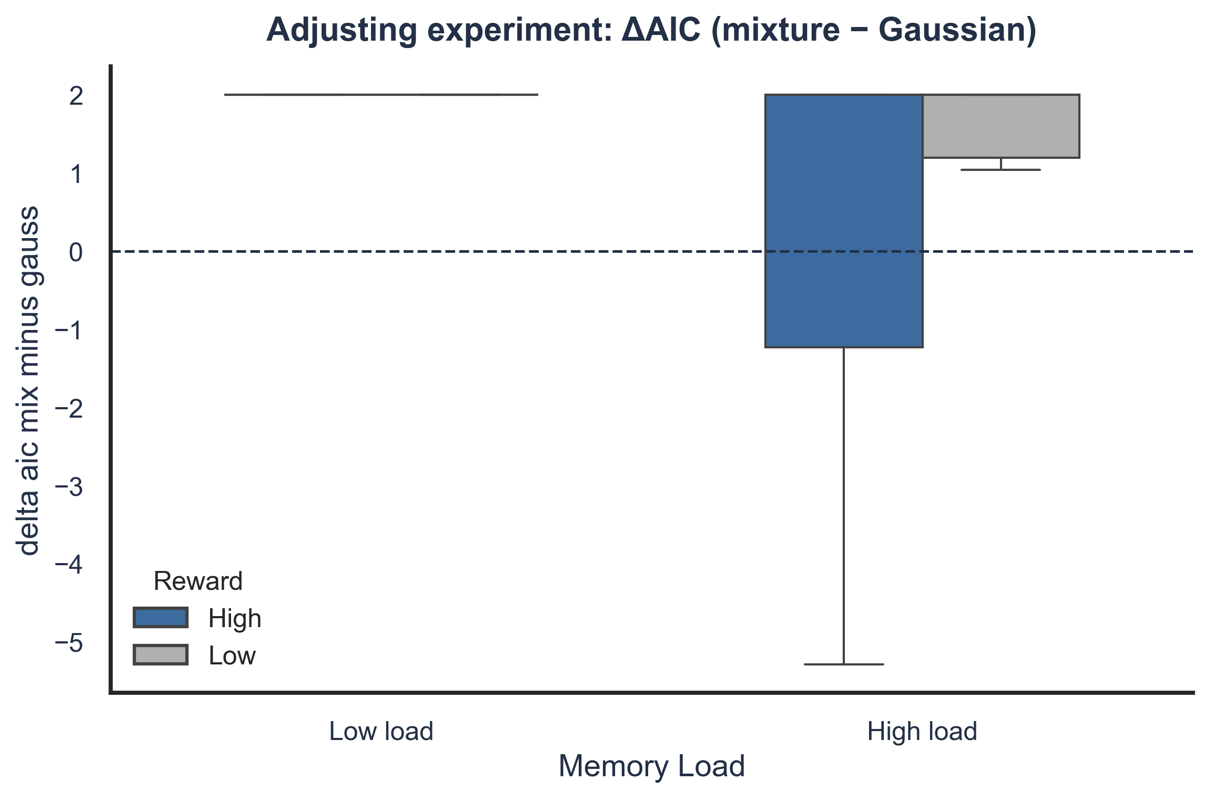

**A**

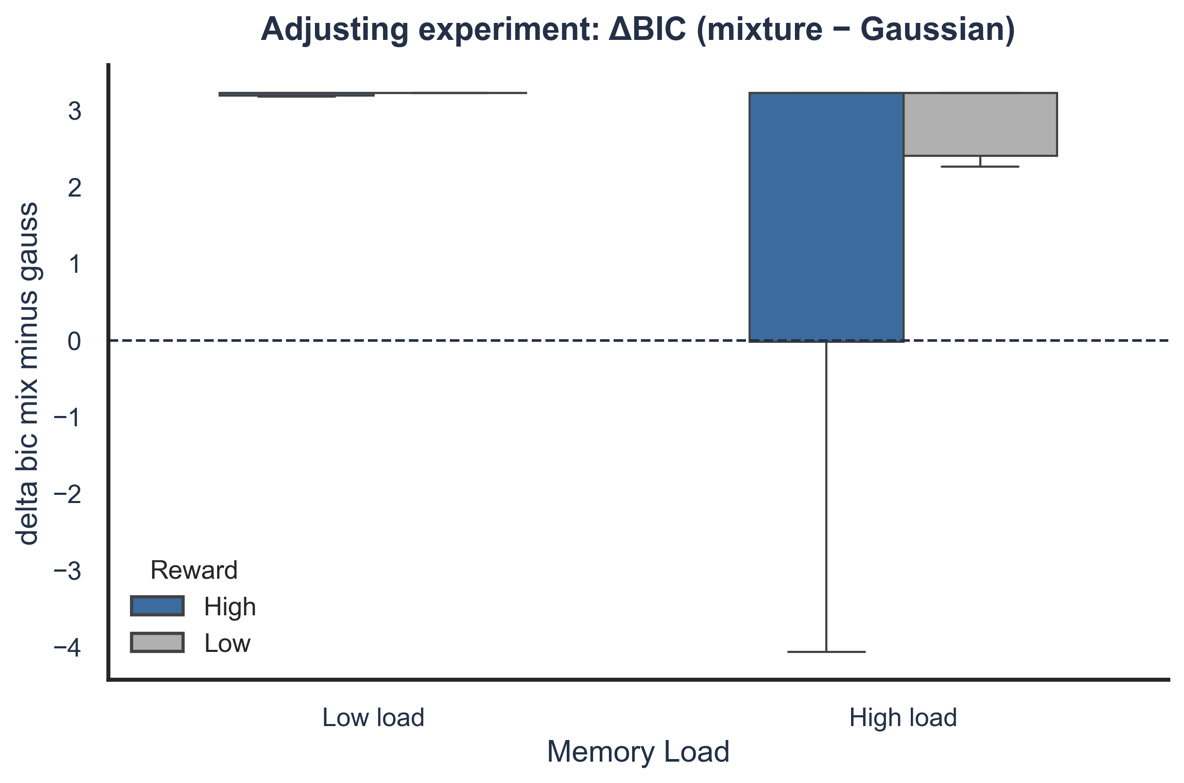

**C**

**B**

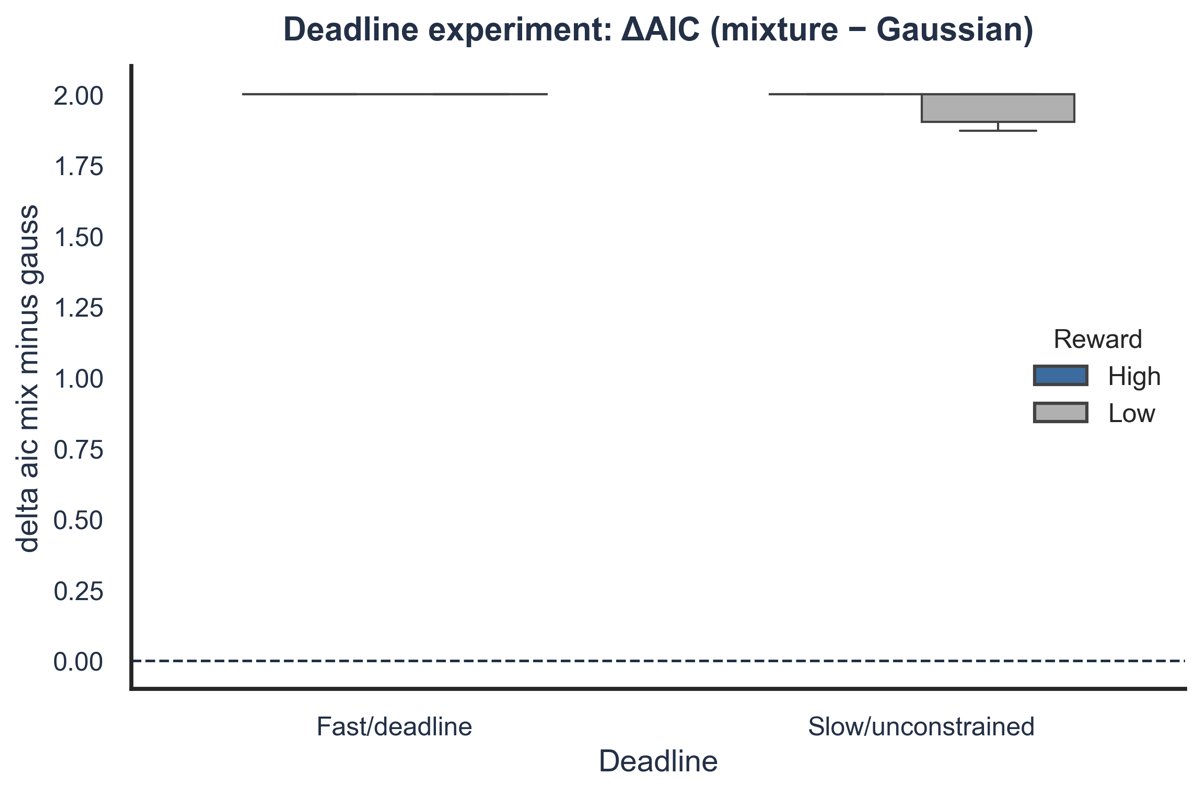

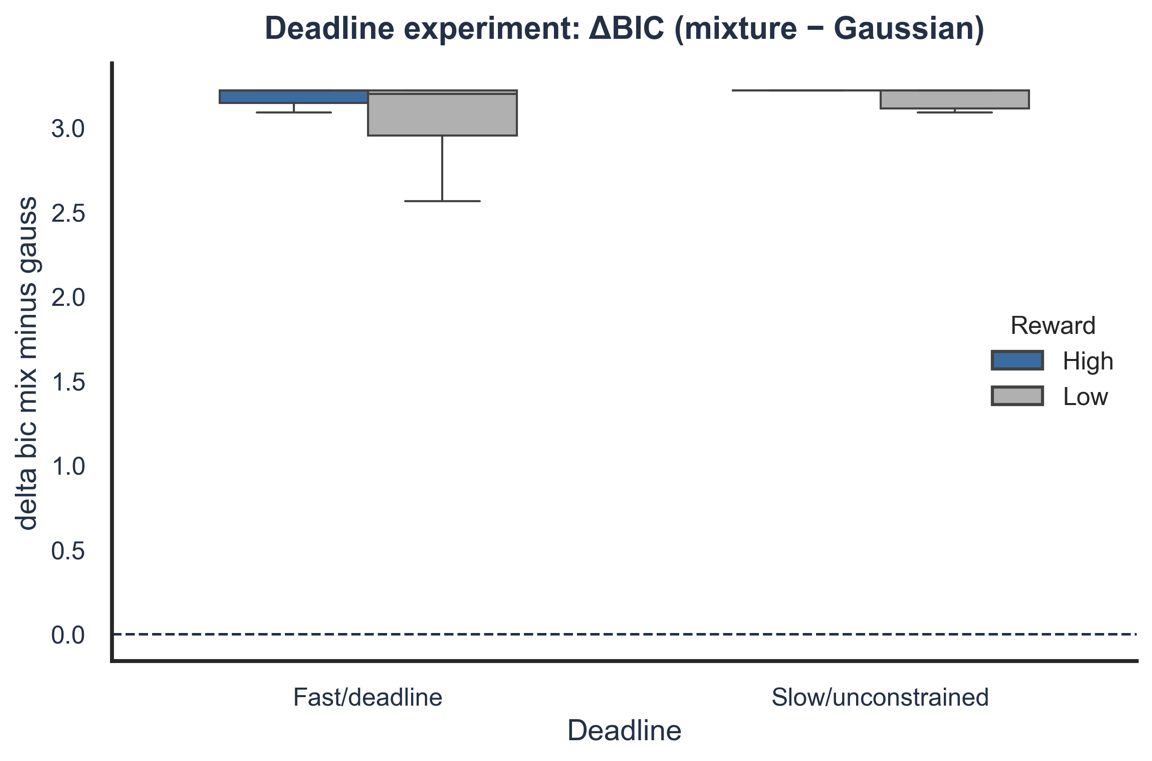

**D**

**Fig.S2. Model comparison metrics for the Gaussian-uniform mixture model versus the Gaussian-only model.** Distributions of the difference in information criteria between the mixture model and the Gaussian-only model, computed separately for each participant and condition. Positive values favour the Gaussian-only model. (**a**) ΔAIC in Experiment 1. (**b**) ΔBIC in Experiment 1. (**c**) ΔAIC in Experiment 2. (**d**) ΔBIC in Experiment 2. Differences were small and inconsistent across participants and conditions (Exp 1: ΔAIC = −0.72 to 1.00; ΔBIC = 0.49 to 2.21; Exp 2: ΔAIC = −0.01 to 1.35; ΔBIC = 1.21 to 2.57), and the Gaussian-only model was preferred more often than the mixture model, indicating that behaviour was well-described by a single memory-based process.

**A**

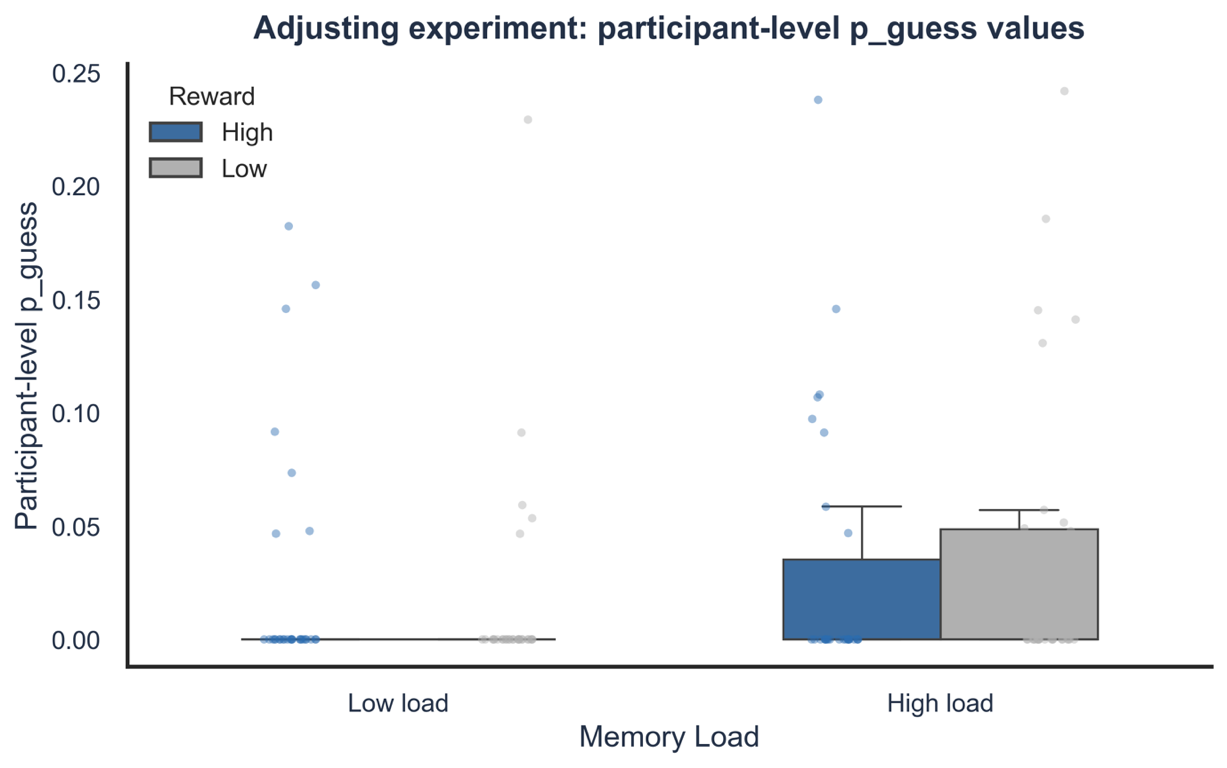

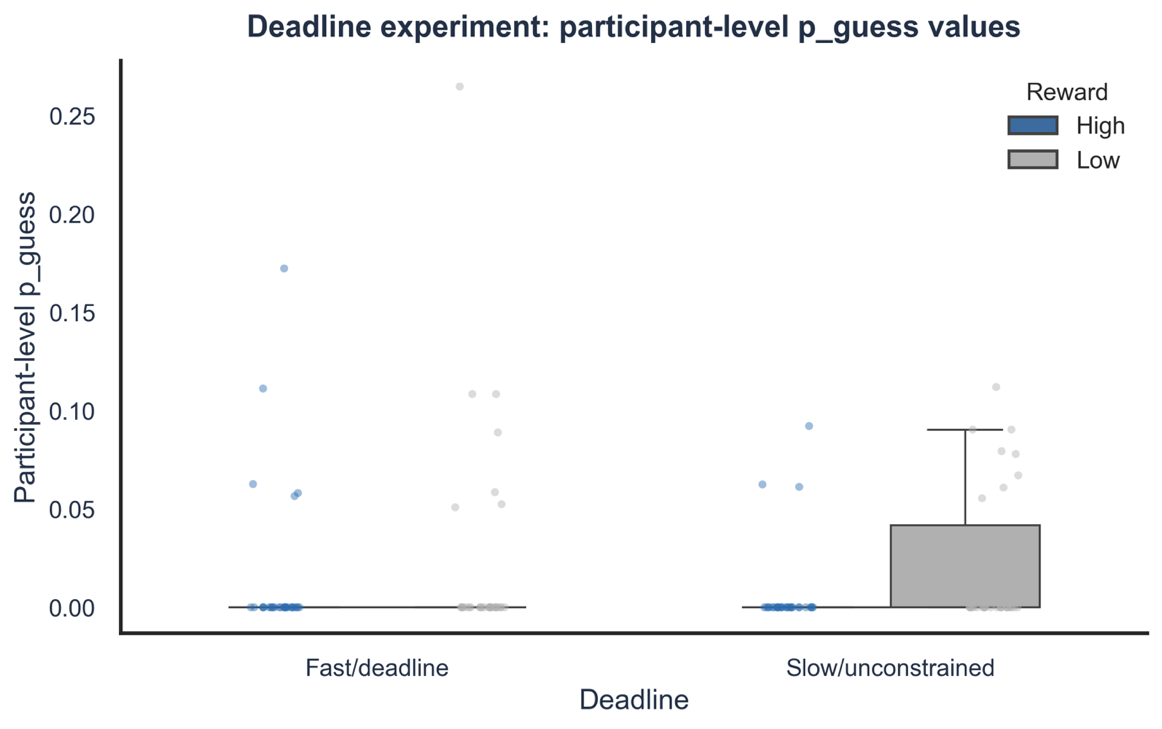

**B**

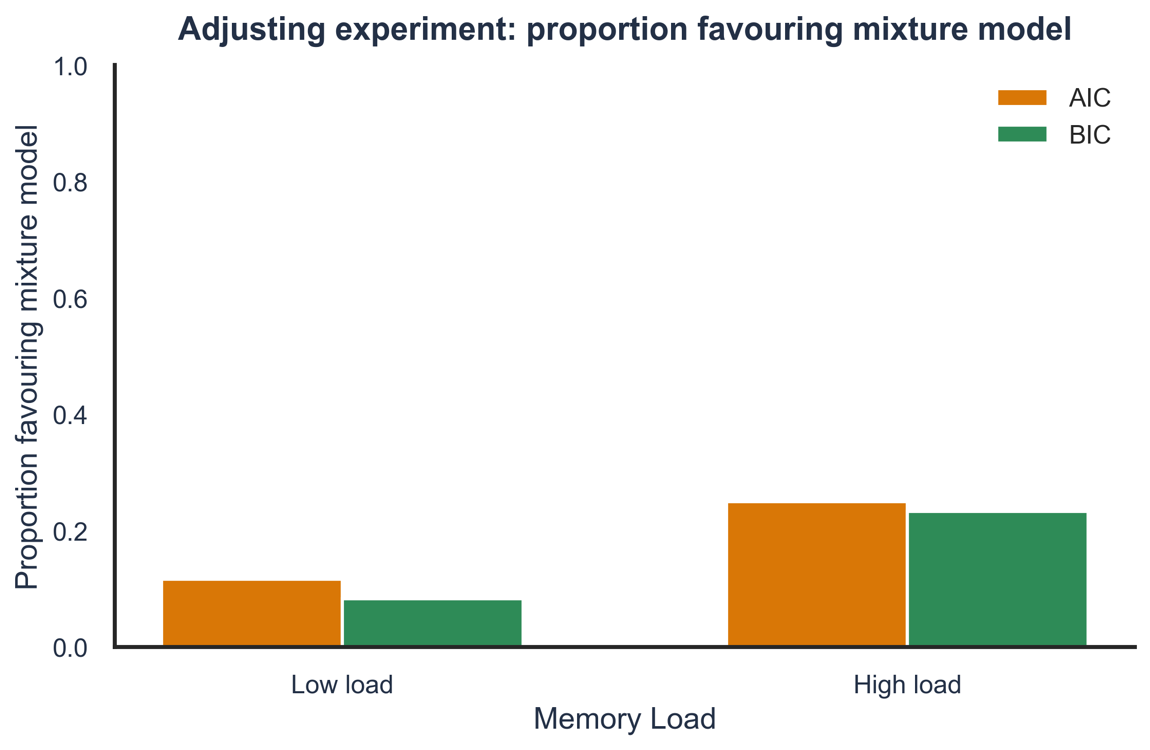

**C**

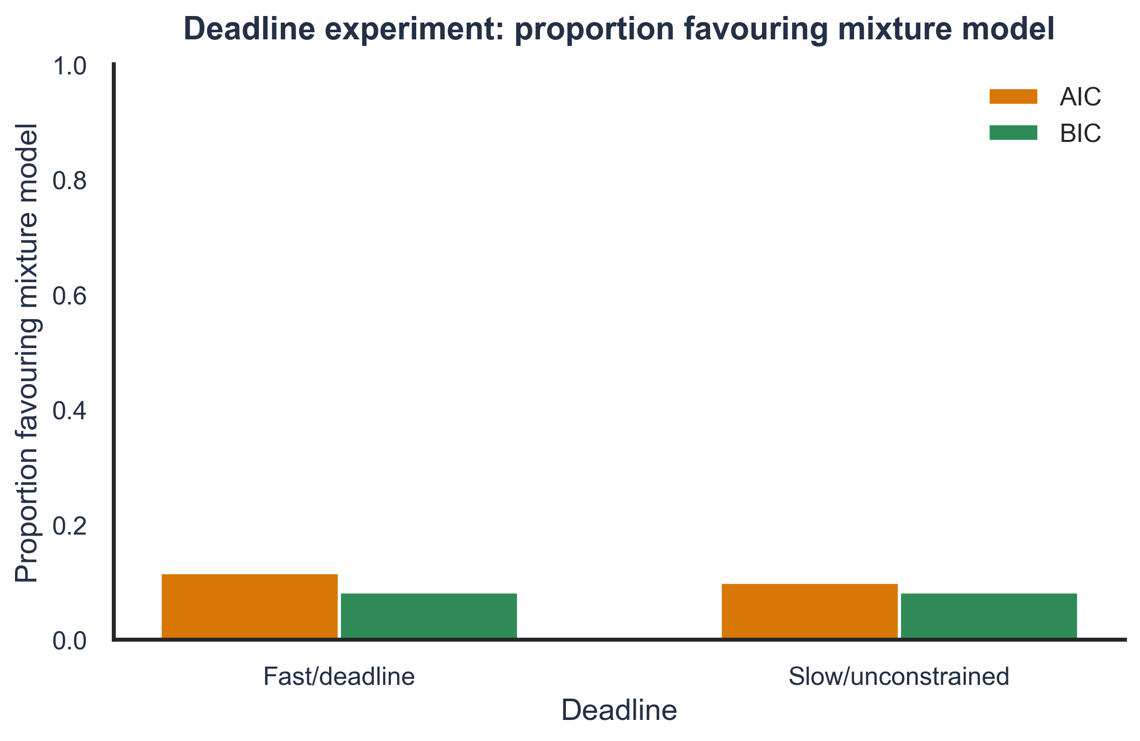

**D**

**Fig.S3. Distribution of p_guess estimates and model preference across experiments.** Top row: distribution of participant-level p_guess estimates from the Gaussian-uniform mixture model. (**a**) Experiment 1. (**b**) Experiment 2. Most participants (~75–80%) had p_guess estimates near zero, indicating minimal random guessing. Bottom row: proportion of participants for whom the Gaussian-uniform mixture model was preferred over the Gaussian-only model, based on the Akaike Information Criterion (AIC) and Bayesian Information Criterion (BIC). (**c**) Experiment 1. (**d**) Experiment 2. The mixture model was preferred in only a minority of participants and conditions, further supporting the conclusion that behaviour was well-described by a single memory-based process.

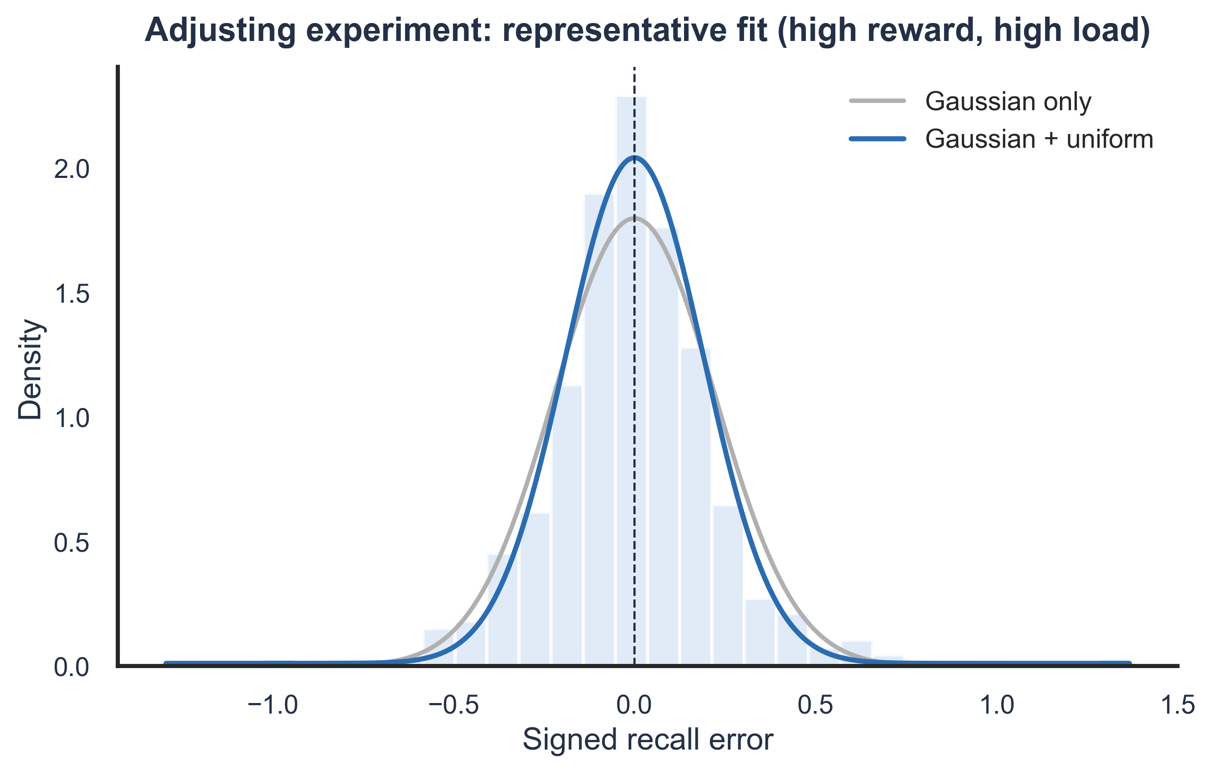

**A**

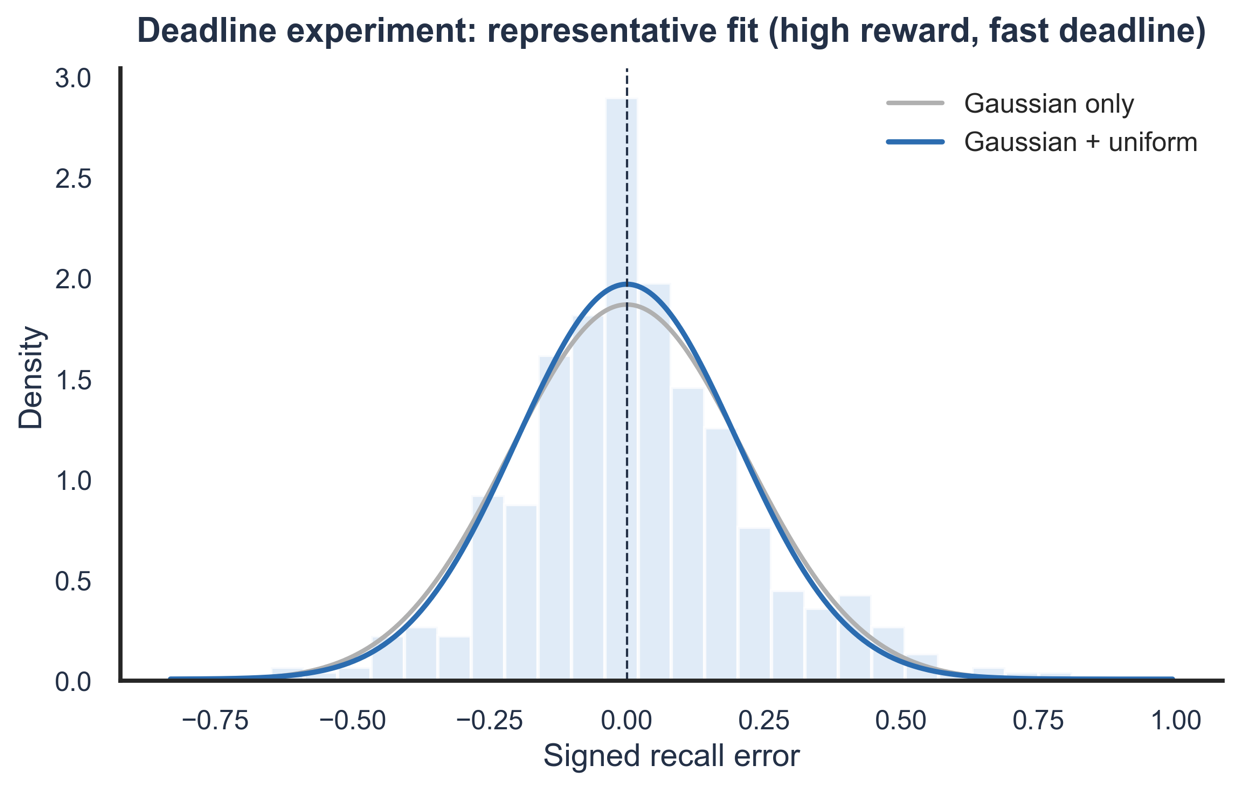

**B**

**Fig.S4. Representative mixture-model fits to signed recall error distributions.** Empirical distributions of signed recall error (grey histograms) overlaid with fitted Gaussian-uniform mixture model densities (solid curves) for a representative participant. (**a**) Experiment 1. (**b**) Experiment 2. The model captured the empirical error distributions well, with responses concentrated around zero and minimal contribution from the uniform (guessing) component.

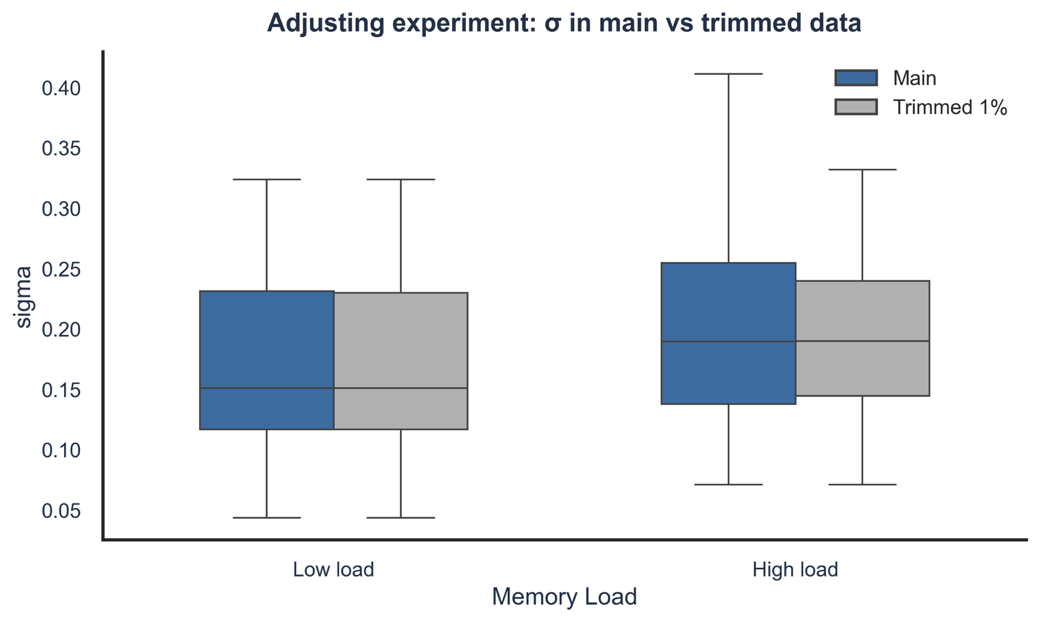

**A**

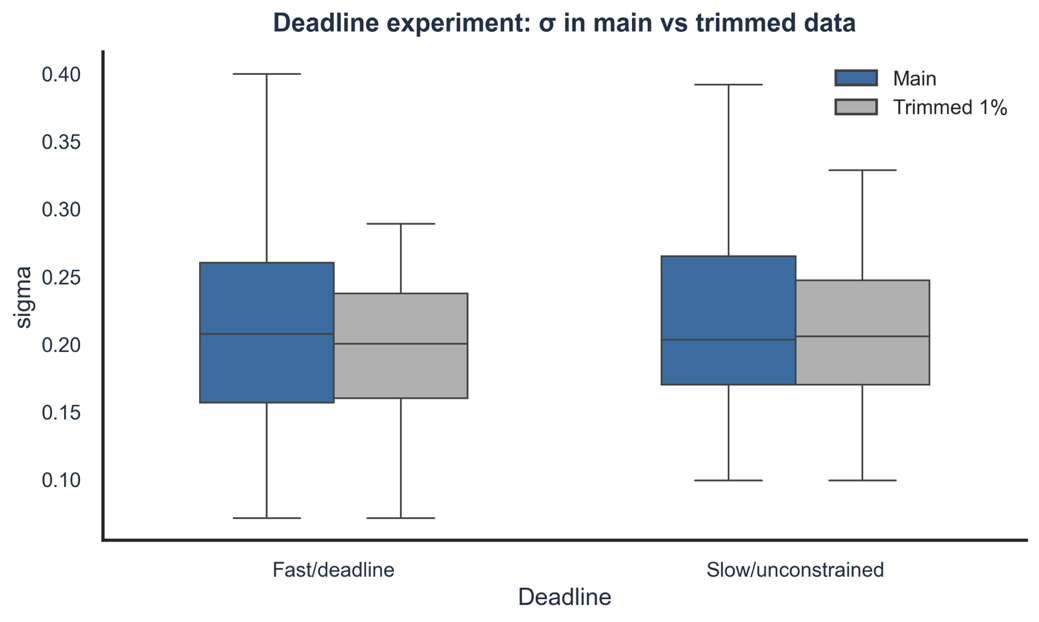

**B**

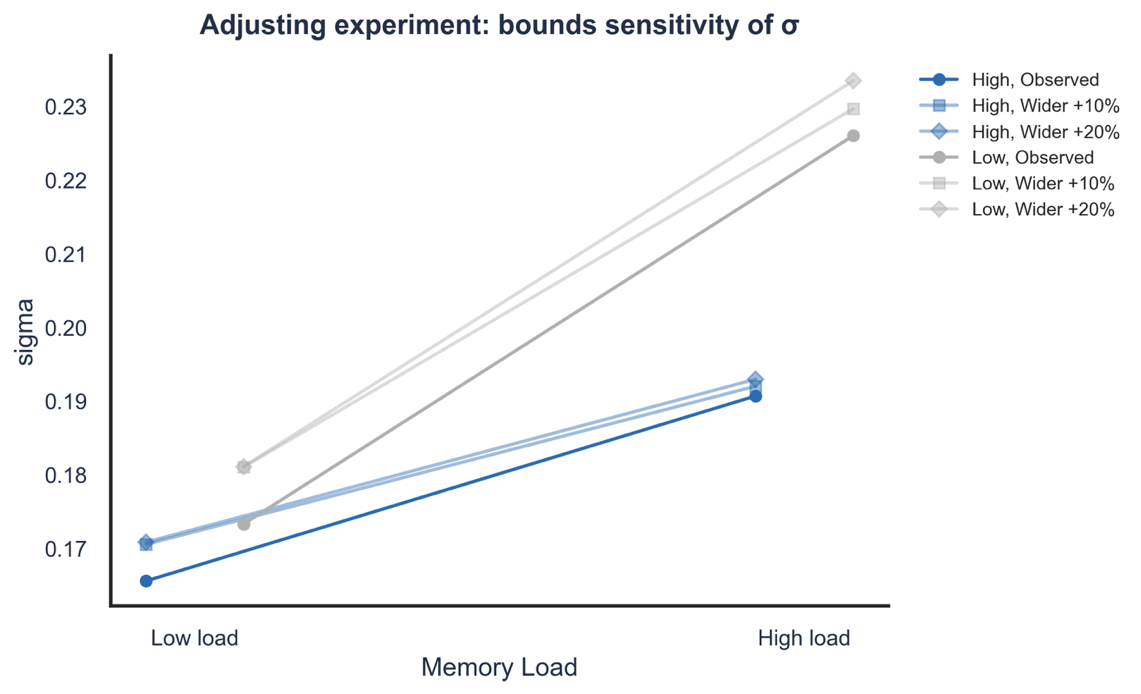

**C**

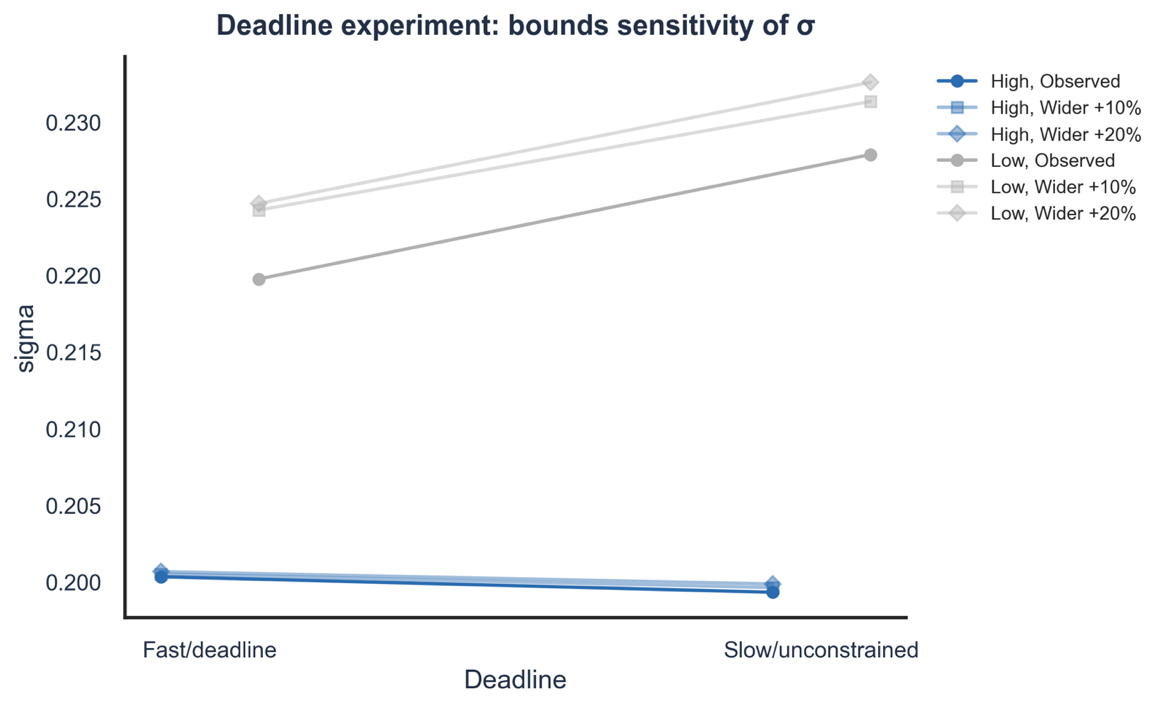

**D**

**Fig.S5. Robustness of σ estimates to trimming and parameter bounds.** Top row: effect of excluding extreme responses (top and bottom 1% of trials) on σ estimates. (**a**) Experiment 1 (Reward × Memory Load). (**b**) Experiment 2 (Reward × Deadline). Bottom row: effect of varying the assumed bounds of the uniform distribution (±10–20% of the observed error range) on σ estimates. (**c**) Experiment 1. (**d**) Experiment 2. Reward-related reductions in σ remained consistent across all specifications, indicating robustness to modelling assumptions and extreme responses. Error bars show standard errors of the mean.

**B**

**A**

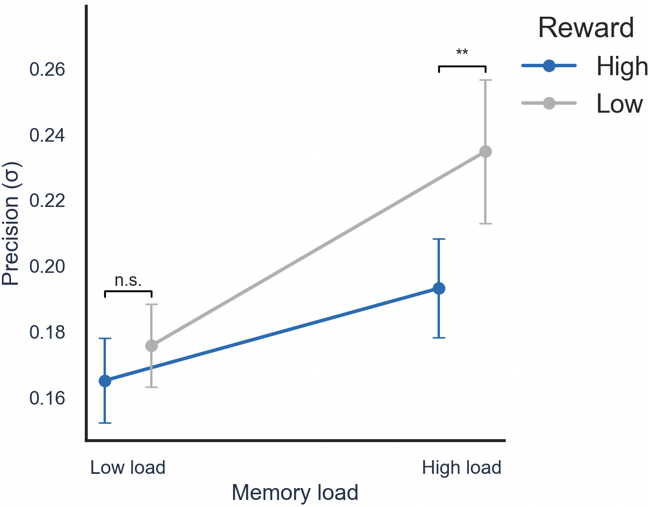

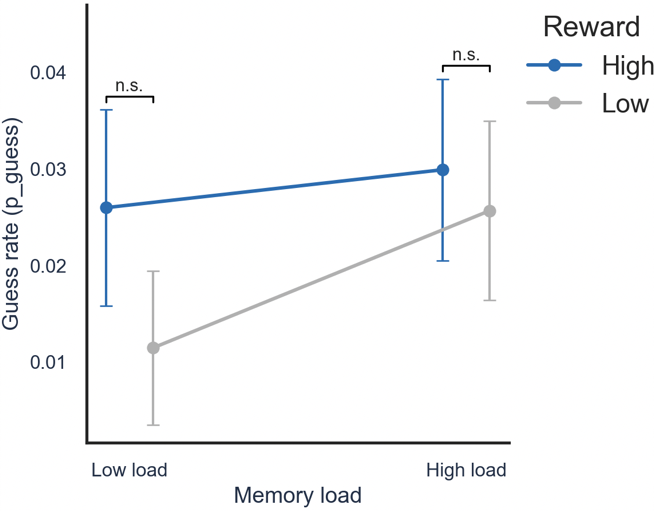

**Fig.S6. Reward effect on mixture-model parameters on non-resubmitted trials only (Experiment 1).** Main mixture-model analysis re-run using only trials on which participants did not resubmit (2,589 of 2,983 trials; 86.8%). (**a**) Precision parameter σ (units: seconds). Reward significantly reduced σ, F(1,29) = 7.70, P = .010, BF₁₀ = 4.68, comparable in magnitude and evidence to the main analysis (**Fig.5a**). (**b**) Guess rate p_guess. Reward did not affect p_guess, F(1,29) = 1.03, P = .319, BF₀₁ = 3.22, consistent with the main analysis (**Fig.5b**). Error bars show standard errors of the mean.

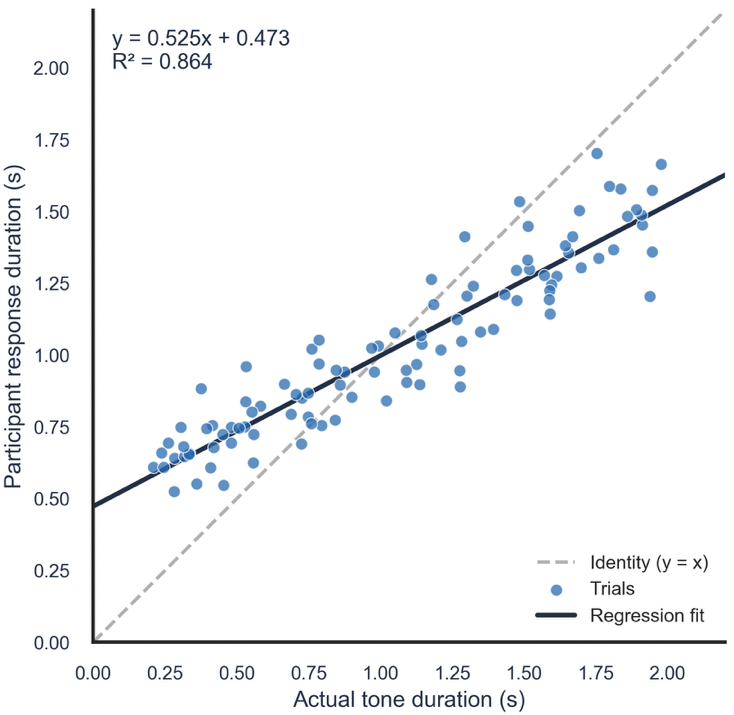

**Fig. S7. Schematic illustration of the participant-level linear regression used to compute absolute recall error.** Simulated data demonstrating the analytical approach, showing the relationship between target tone duration (x-axis) and reproduced duration (y-axis), with the fitted linear regression line. The simulated pattern reproduces the central tendency bias observed empirically (Vierordt's law): shorter tones tend to be overestimated and longer tones underestimated. In the actual analysis, absolute recall error was defined as the absolute residual from this participant-level regression fitted separately to each participant's real data, providing a measure of deviation from the expected response independent of central tendency bias.
